# Reward-based training of recurrent neural networks for cognitive and value-based tasks

**DOI:** 10.1101/070375

**Authors:** H. Francis Song, Guangyu R. Yang, Xiao-Jing Wang

## Abstract

Trained neural network models, which exhibit many features observed in neural recordings from behaving animals and whose activity and connectivity can be fully analyzed, may provide insights into neural mechanisms. In contrast to commonly used methods for supervised learning from graded error signals, however, animals learn from reward feedback on definite actions through reinforcement learning. Reward maximization is particularly relevant when the optimal behavior depends on an animal’s internal judgment of confidence or subjective preferences. Here, we describe reward-based training of recurrent neural networks in which a value network guides learning by using the selected actions and activity of the policy network to predict future reward. We show that such models capture both behavioral and electrophysiological findings from well-known experimental paradigms. Our results provide a unified framework for investigating diverse cognitive and value-based computations, including a role for value representation that is essential for learning, but not executing, a task.

## Introduction

A major challenge in uncovering the neural mechanisms underlying complex behavior is our incomplete access to relevant circuits in the brain. Recent work has shown that model neural networks optimized for a wide range of tasks, including visual object recognition (Cadieu et al., 2014; Yamins et al., 2014; Hong et al., 2016), perceptual decision-making and working memory (Mante et al., 2013; Barak et al., 2013; Carnevale et al., 2015; Song et al., 2016; Miconi, 2016), timing and sequence generation (Laje & Buonomano, 2013; Rajan et al., 2015), and motor reach (Hennequin et al., 2014; Sussillo et al., 2015), can reproduce important features of neural activity recorded in numerous cortical areas of behaving animals. The analysis of such circuits, whose activity and connectivity are fully known, has therefore re-emerged as a promising tool for understanding neural computation (Zipser & Andersen, 1988; Sussillo, 2014; Gao & Ganguli, 2015). Constraining network training with tasks for which detailed neural recordings are available may also provide insights into the principles that govern learning in biological circuits (Sussillo et al., 2015; Song et al., 2016).

Previous applications of this approach to “cognitive-type” behavior such as perceptual decision-making and working memory have focused on supervised learning from graded error signals. Animals, however, learn to perform specific tasks from reward feedback provided by the experimentalist in response to definite actions, i.e through reinforcement learning (Sutton & Barto, 1998). Unlike in supervised learning where the network must be given the correct response on each trial, reinforcement learning merely provides evaluative feedback to the network, i.e whether the selected action was good or bad. For the purposes of using model neural networks to generate hypotheses about neural mechanisms, this is particularly relevant in tasks where the optimal behavior depends on an animal’s internal state or subjective preferences. In a perceptual decision-making task with postdecision wagering, for example, on some trials the animal can opt for a “sure” choice that results in a small (compared to the correct choice) but certain reward (Kiani & Shadlen, 2009). The optimal decision regarding whether or not to select the sure choice depends not only on the task condition, such as the proportion of coherently moving dots, but also on the animal’s own confidence in its decision *during the trial*. Learning to make this judgment cannot be reduced to minimizing a trial-by-trial error signal; it can be learned, however, by choosing the actions that result in greatest reward. Meanwhile, supervised learning is inherently unsuitable for value-based, or economic, decision-making where the “correct” judgment depends explicitly on rewards associated with different actions, even for identical sensory inputs (Padoa-Schioppa & Assad, 2006). More fundamentally, reward plays a central role in all types of animal learning (Sugrue et al., 2005). Explicitly incorporating reward into network training is therefore a necessary step toward elucidating the biological substrates of learning, in particular reward-dependent synaptic plasticity (Seung, 2003; Soltani et al., 2006; Izhikevich, 2007; Urbanczik & Senn, 2009; Frémaux et al., 2010; Soltani & Wang, 2010; Hoerzer et al., 2014; Brosch et al., 2015; Friedrich & Lengyel, 2016) and the role of different brain structures in learning (Frank & Claus, 2006).

In this work, we build on advances in policy gradient reinforcement learning, specifically the REINFORCE algorithm (Williams, 1992; Sutton et al., 2000; Peters & Schaal, 2008;Wierstra et al., 2009), to demonstrate reward-based training of recurrent neural networks (RNNs) for several well-known experimental paradigms in systems neuroscience. The networks consist of two modules in an “actor-critic” architecture (Barto et al., 1983; Grondman et al., 2012), in which a policy network uses inputs provided by the environment to select actions that maximize reward, while a value network uses the selected actions and activity of the policy network to predict future reward and guide learning. We first present networks trained for tasks that have been studied previously using various forms of supervised learning (Mante et al., 2013; Barak et al., 2013; Song et al., 2016); they are characterized by “simple” input-output mappings in which the correct response for each trial depends only on the task condition, and include perceptual decision-making, context-dependent integration, multisensory integration, and parametric working memory tasks. We then show results for tasks in which the optimal behavior depends on the animal’s internal judgment of confidence or subjective preferences, specifically a perceptual decision-making task with postdecision wagering (Kiani & Shadlen, 2009) and a value-based economic choice task (Padoa-Schioppa & Assad, 2006). Interestingly, unlike for the other tasks where we focus on comparing the activity of units in the policy network to neural recordings in the prefrontal and posterior parietal cortex of animals performing the same tasks, for the economic choice task the activity of the value network exhibits a striking resemblance to neural recordings from the orbitofrontal cortex (OFC), which has long been implicated in the representation of reward-related signals (Wallis, 2007).

Indeed, a crucial feature of our REINFORCE-based model is that a reward baseline—in this case a recurrently connected value network—is essential for learning, but not for executing the task, because the latter depends only on the policy network. Importantly, learning can still occur without the value network but is much more difficult. It is sometimes observed in experiments that reward-modulated structures in the brain such as the basal ganglia or orbitofrontal cortex are necessary for learning or adapting to a changing environment, but not for executing a previously learned skill (Turner & Desmurget, 2010; Schoenbaum et al., 2011; Stalnaker et al., 2015). This suggests that one possible role for such circuits may be representing an accurate baseline to guide learning. Moreover, since confidence is closely related to expected reward in many cognitive tasks, the explicit computation of expected reward by the value network provides a concrete, learning-based rationale for confidence estimation as a fundamental and ubiquitous component of decision-making (Kepecs et al., 2008; Wei & Wang, 2015), even when it is not strictly required for performing the task.

Conceptually, the formulation of behavioral tasks in the language of reinforcement learning presented here is closely related to the solution of partially observable Markov decision processes (POMDPs) (Kaelbling et al., 1998) using either model-based belief states (Rao, 2010) or model-free working memory (Todd et al., 2008). Indeed, as in Dayan & Daw (2008) one of the goals of this work is to unify related computations into a common language that is applicable to a wide range of tasks in systems neuroscience. However, the formulation using policies represented by RNNs allows for a far more general description, and, in particular, makes the assumption of a Markovian environment unnecessary (Wierstra et al., 2009). Such policies can also be compared more directly to “optimal” solutions when they are known, for instance to the signal detection theory account of perceptual decision-making (Gold & Shadlen, 2007). Thus, in addition to expanding the range of tasks and neural mechanisms that can be studied with trained RNNs, our work provides a convenient framework for the study of cognitive and value-based computations in the brain, which have often been viewed from distinct perspectives but in fact arise from the same reinforcement learning paradigm.

## Results

### Policy gradient reinforcement learning for behavioral tasks

For concreteness, we illustrate the following in the context of a simplified perceptual decision-making task based on the random dots motion (RDM) discrimination task as described in Kiani et al. (2008) (Figure 1A). In its simplest form, in an RDM task the monkey must maintain fixation until a “go” cue indicates that the monkey should make a decision regarding the direction of coherently moving dots on the screen. Thus the three possible actions available to the monkey at any given time are fixate, choose left, or choose right. The true direction of motion, which can be considered a *state* of the environment, is not known to the monkey with certainty, i.e is *partially observable*. The monkey must therefore use the noisy sensory evidence to infer the direction in order to select the correct response at the end of the trial. Breaking fixation early results in a negative reward in the form of a timeout, while giving the correct response after the fixation cue is extinguished results in a positive reward in the form of juice. The goal of this section is to give a general description of such tasks and how an animal or RNN can learn a behavioral *policy* for choosing actions at each time to maximize the reward.

**Figure 1.**
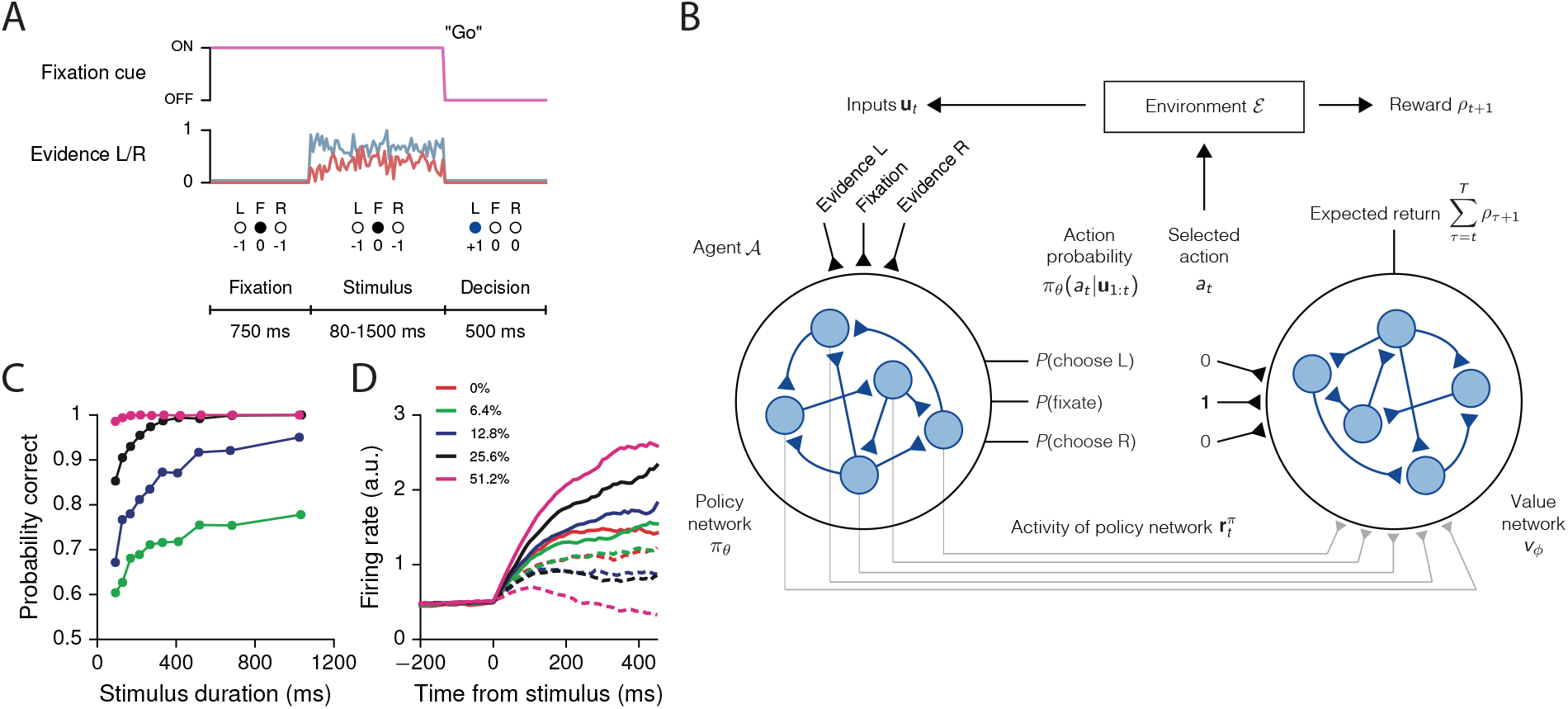
Recurrent neural networks for reinforcement learning. (**A**) Task structure for a simple perceptual decision-making task with variable stimulus duration. The agent must maintain fixation (*a*_*t*_=F) until the go cue, which indicates the start of a decision period during which choosing the correct response (*a*_*t*_=L or *a*_*t*_=R) results in a positive reward. The agent receives zero reward for responding incorrectly, while breaking fixation early results in an aborted trial and negative reward. (**B**) At each time *t* the agent selects action *a*_*t*_ according to the output of the policy network *π*_*θ*_, which can depend on all past and current inputs **u**_1:*t*_ provided by the environment. In response, the environment transitions to a new state and provides reward *ρ*_*t*+1_ to the agent. The value network 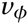 uses the selected action and the activity of the policy network 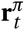 to predict future rewards. All the weights shown are plastic, i.e trained by gradient descent. (**C**) Performance of the network trained for the task in (A), showing the percent correct by stimulus duration, for different coherences (the difference in strength of evidence for L and R). (**D**) Neural activity of an example policy network unit, sorted by coherence and aligned to the time of stimulus onset. Solid lines are for positive coherence, dashed for negative coherence. Figure supplement 1. Learning curves for the simple perceptual decision-making task. Figure supplement 2. Output of the value network indicating expected reward. Figure supplement 3. Reaction-time version of the perceptual decision-making task, in which the go cue coincides with the onset of stimulus, allowing the agent to choose when to respond. Figure supplement 4. Learning curves for the reaction-time version of the simple perceptual decisionmaking task. Figure supplement 5. Learning curves for the simple perceptual decision-making task with a linear readout of the policy network as the baseline.

Consider a typical interaction between an experimentalist and animal, which we more generally call the environment *ℰ* and agent *A*, respectively (Figure 1B). At each time *t* the agent chooses to perform actions **a**_*t*_ after observing inputs **u**_*t*_ provided by the environment, and the probability of choosing actions **a**_*t*_ is given by the agent’s policy 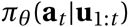 with parameters *θ*. Here the policy is implemented as an RNN, so that *θ* comprises the connection weights, biases, and initial state of the network. The policy at time *t* can depend on all past and current inputs **u**_1:*t*_ = (**u**_1_,**u**_2_, …,**u**_*t*_), allowing the agent to integrate sensory evidence or use working memory to perform the task. The exception is at *t* = 0, when the agent has yet to interact with the environment and selects its actions “spontaneously” according to 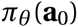. We note that, if the inputs give exact information about the environmental state, i.e., if **u**_*t*_ = **s**_*t*_, then the task becomes a Markov decision process. In general, however, the inputs only provide partial information about the environmental states, requiring the network to accumulate evidence over time to determine the state of the world. In this work we only consider cases where the agent chooses one out of *N*_*a*_ possible actions at each time, so that 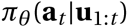 for each *t* is a discrete, normalized probability distribution over the possible actions *a*_1_, …,*a*_*N*__*a*_. More generally, **a**_*t*_ can implement several distinct actions or even continuous actions by representing, for example, the means of Gaussian distributions (Peters & Schaal, 2008; Wierstra et al., 2009). After each set of actions by the agent at time *t* the environment provides a reward (or special observable) *ρ*_*t* + 1_ at time *t*+1, which the agent attempts to maximize in the sense described below.

In the case of the example RDM task above (Figure 1A), the environment provides (and the agent receives) as inputs a fixation cue and noisy evidence for two choices L and R during a variable-length stimulus presentation period. The strength of evidence, or the difference between the evidence for L and R, is called the coherence, and in the actual RDM experiment corresponds to the percentage of dots moving coherently in one direction on the screen. The agent chooses to performone of *N*_*a*_ = 3 actions at each time: fixate (*a*_*t*_ = F), choose L (*a*_*t*_ = L), or choose R (*a*_*t*_ = R). In this task, the agent must choose F as long as the fixation cue is on, and then, when the fixation cue is turned off to indicate that the agent should make a decision, correctly choose L or R depending on the sensory evidence. Indeed, for all tasks in this work we required that the network “make a decision” (i.e., break fixation to indicate a choice at the appropriate time) on at least 99% of the trials, whether the response was correct or not. A trial endswhen the agent chooses L or R regardless of the task epoch: breaking fixation early before the go cue results in an aborted trial and a negative reward *ρ*_*t*_ = − 1, while a correct decision is rewarded with *ρ*_*t*_ = + 1. Making the wrong decision results in no reward, *ρ*_*t*_ = 0. For the zero-coherence condition the agent is rewarded randomly on half the trials regardless of its choice. Otherwise the reward is always *ρ*_*t*_ = 0.

Formally, a trial proceeds as follows. At time *t* = 0, the environment is in state **s**_0_ with probability ℰ (**s**_0_). The state **s**_0_ can be considered the starting time (i.e., *t* = 0) and “task condition,” which in the RDM example consists of the direction of motion of the dots (i.e., whether the correct response is L or R) and the coherence of the dots (the difference between evidence for L and R). The time component of the state, which is updated at each step, allows the environment to present different inputs to the agent depending on the task epoch. The true state **s**_0_ (such as the direction of the dots) is only partially observable to the agent, so that the agent must instead infer the state through inputs **u**_*t*_ provided by the environment during the course of the trial. As noted previously, the agent initially chooses actions **a**_0_ with probability 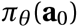. The networks in this work almost always begin by choosing F, or fixation.

At time *t* = 1, the environment, depending on its previous state **s**_0_ and the agent’s action **a**_0_, transitions to state **s**_1_ with probability ℰ (**s**_1_|**s**_0_,**a**_0_) and generates reward *ρ*_1_. In the perceptual decision-making example, only the time advances since the trial condition remains constant throughout. From this state the environment generates observable **u**_1_ with a distribution given by 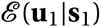. If *t* = 1 were in the stimulus presentation period, for example, **u**_1_ would provide noisy evidence for L or R, as well as the fixation cue. In response, the agent, depending on the inputs **u**_1_ it receives from the environment, chooses actions **a**_1_ with probability 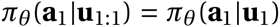. The environment, depending on its previous states **s**_0:1_ = (**s**_0_,**s**_1_) and the agent’s previous actions *a*_0:1_ = (**a**_0_,**a**_1_), then transitions to state **s**_2_ with probability E (**s**_2_|**s**_0:1_,**a**_0:1_) and generates reward *ρ*_2_. These steps are repeated until the end of the trial at time *T*. Trials can terminate before *T* (e.g., for breaking fixation early), so that *T* represents the maximum length of a trial. In order to emphasize that rewards follow actions, we adopt the convention in which the agent performs actions at *t* = 0, …,*T* and receives rewards at *t* = 1, …,*T* +1.

The goal of the agent is to maximize the sum of expected future rewards at time *t* = 0, or expected return

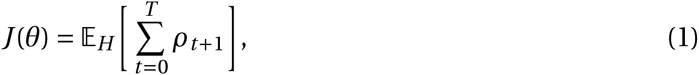

where the expectation 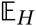 is taken over all possible trial histories 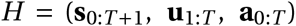 consisting of the states of the environment, the inputs given to the agent, and the actions of the agent. In practice, the expectation value in Equation 1 is estimated by performing *N*_trials_ trials for each policy update, i.e., with a Monte Carlo approximation. Here we only consider the episodic, “finite-horizon” case where *T* is finite. The expected return depends on the policy and hence parameters *θ*, and we use Adam stochastic gradient descent (SGD) (Kingma & Ba, 2015) with gradient clipping (Graves, 2013; Pascanu et al., 2013b) to find the parameters that maximize this reward (Methods).

More specifically, for the policy network we use gradient descent to minimize an objective function 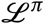of the form

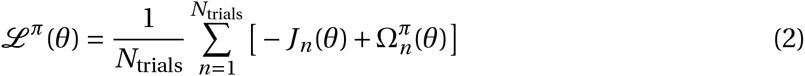

with respect to the connection weights, biases, and initial state of the policy network,which we collectively denote as *θ*. Here 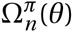 contains any regularization terms for the policy network, for instance an entropy term to control the degree of exploration (Xu et al., 2015). The key gradient 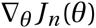 is given for each trial *n* by the REINFORCE algorithm (Williams, 1992; Sutton et al., 2000; Peters & Schaal, 2008;Wierstra et al., 2009) as

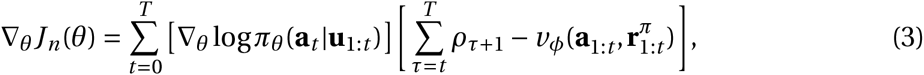

where 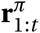 are the firing rates of the policy network units up to time *t*, 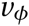 denotes the value function as described below, and the gradient 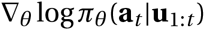, known as the *eligibility*, [and likewise 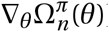] is computed by backpropagation through time (BPTT) (Rumelhart et al., 1986) for the selected actions **a**_*t*_. The sum over rewards in large brackets only runs over 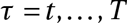, which reflects the fact that actions do not affect past rewards. In this form the terms in the gradient have the intuitive property that they are nonzero only if the actual return deviates from what was predicted by the baseline. This reward baseline is an important feature in the success of almost all REINFORCE-based algorithms, and is here represented by a second RNN 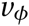 with parameters 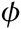 in addition to the policy network 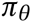. This baseline network, which we call the *value network*, uses the selected actions **a**_1:*t*_ and activity of the policy network 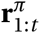 to predict the expected return at each time *t* = 1, …,*T* (the value network also predicts the expected return at *t* = 0 based on its own initial states, with the understanding that 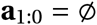 and 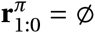 are empty sets). The value network is trained by minimizing a second objective function

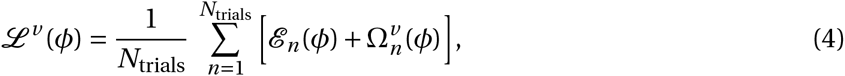

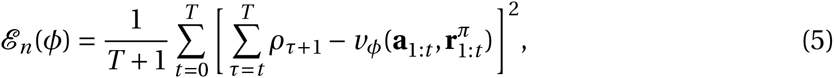

where 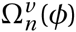 denotes any regularization terms for the value network. The necessary gradient 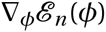 [and likewise 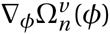] is again computed by BPTT.

### Policy and value recurrent neural networks

The policy probability distribution 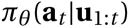 and scalar baseline 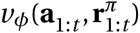 are each represented by an RNN of *N* firing-rate units **r**^*π*^and **r**^*v*^, respectively, where we interpret each unit as the mean firing rate of a group of neurons. In the case where the agent chooses a single action at each time *t*, the activity of the policy network determines 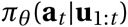 through a linear readout followed by softmax normalization:

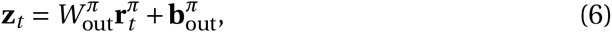

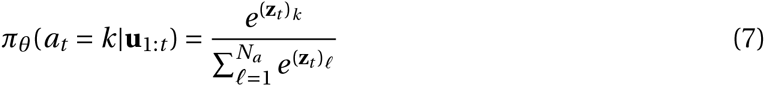

for *k* = 1, …,*N*_*a*_. Here 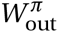 is an *N*_*a*_×*N* matrix of connection weights from the units of the policy network to the *N*_*a*_ linear readouts **z**_*t*_, and 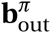 are *N*_*a*_ biases. Action selection is implemented by randomly sampling from the probability distribution in Equation 7, and constitutes an important difference from previous approaches to training RNNs for cognitive tasks, namely, here the final output of the network (during training) is a specific action, not a graded decision variable. From a biological point of view, we consider this sampling as an abstract representation of the downstream action selection mechanisms present in the brain, including the role of noise in implicitly realizing stochastic choices with deterministic outputs (Wang, 2002, 2008). Meanwhile, the activity of the value network predicts future returns through a linear readout

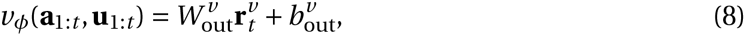

where 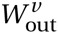 is an 1×*N* matrix of connection weights from the units of the value network to the single linear readout 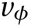, and 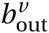 is a bias term.

In order to take advantage of recent developments in training RNNs [in particular, addressing the problem of vanishing gradients (Bengio et al., 1994)] while retaining intepretability, we use a modified form of Gated Recurrent Units (GRUs) (Cho et al., 2014; Chung et al., 2014) with a threshold-linear “*f* -*I* ” curve [*x*]_+_ = max(0,*x*) to obtain positive firing rates. These units are thus leaky, threshold-linear units with dynamic time constants and gated recurrent inputs. The equations that describe their dynamics can be derived by a naïve discretization of the following continuous-time equations for the *N* currents **x** and corresponding rectified linear firing rates **r**:

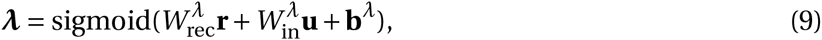

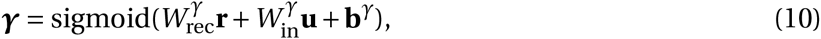

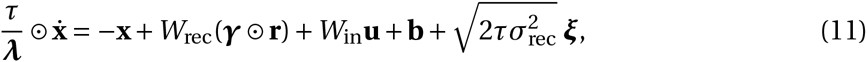

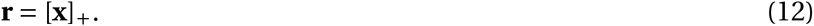

Here 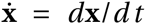 is the derivative of **x** with respect to time, 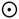 denotes elementwise multiplication, sigmoid(*x*) = [1+*e*^−*x*^]^−1^ is the logistic sigmoid, **b**^*λ*^, **b**^*γ*^, and **b** are biases, ***ξ*** are *N* independent Gaussian white noise processes with zero mean and unit variance, and 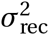 controls the size of this noise. The multiplicative gates ***λ*** dynamically modulate the overall time constant *τ* for network units, while the ***γ*** control the recurrent inputs. The *N*×*N* matrices *W*_rec_, 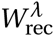, and 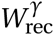 are the recurrent weight matrices, while the *N* ×*N*_in_ matrices *W*_in_, 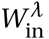*¸* and 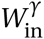 in are connection weights from the *N*_in_ inputs **u** to the *N* units of the network. We note that in the case where ***λ***→1 and ***γ***→1 the equations reduce to “simple” leaky threshold-linear units without the modulation of the time constants or gating of inputs. We constrain the recurrent connection weights (Song et al., 2016) so that the overall connection probability is *p*_*c*_; specifically, the number of incoming connections for each unit, or in-degree, was set to exactly *K* = *p*_*c*_*N*.

The result of discretizing Equations 9-12, as well as details on initializing the network parameters, are given in Methods. We successfully trained networks with time steps Δ*t* = 1 ms, but for computational convenience all of the networks in this work were trained and run with Δ*t* = 10 ms. We note that, for typical tasks in systems neuroscience lasting on the order of several seconds, this already implies trials lasting hundreds of time steps. Unless noted otherwise in the text, all networks were trained using the parameters listed in Table 1.

**Table 1.** Parameters for reward-based recurrent neural network training. Unless noted otherwise in the text, networks were trained and run with the parameters listed here.

| Parameter | Symbol | Default value |
| --- | --- | --- |
| Learning rate | $\eta$ | 0.004 |
| Maximum gradient norm | $\Gamma$ | 1 |
| Size of policy/value network | $N$ | 100 |
| Connection probability (policy network) | $p_c^\pi$ | 0.1 |
| Connection probability (value network) | $p_c^\nu$ | 1 |
| Time step | $\Delta t$ | 10 ms |
| Unit time constant | $\tau$ | 100 ms |
| Recurrent noise | $\sigma_{\text{rec}}^2$ | 0.01 |
| Initial spectral radius for recurrent weights | $\rho_0$ | 2 |
| Number of trials per gradient update | $N_{\text{trials}}$ | # of task conditions |

While the inputs to the policy network *π*_*θ*_ are determined by the environment, the value network always receives as inputs the activity of the policy network **r**^*π*^, together with information about which actions were actually selected (Figure 1B). The value network serves two purposes: first, the output of the value network is used as the baseline in the REINFORCE gradient, Equation 3, to reduce the variance of the gradient estimate (Peters & Schaal, 2008); second, since policy gradient reinforcement learning does not explicitly use a value function but value information is nevertheless implicitly contained in the policy, the value network serves as an explicit and potentially nonlinear readout of this information. In situations where expected reward is closely related to confidence, this may explain, for example, certain disassociations between perceptual decisions and reports of the associated confidence (Lak et al., 2014).

A reward baseline, which transforms accumulated rewards into a prediction error (Schultz et al., 1997; Bayer & Glimcher, 2005), is essential to many learning schemes, especially those based on REINFORCE. Indeed, it has been suggested that in general this baseline should be not only task-specific but stimulus (task-condition)-specific (Frémaux et al., 2010; Engel et al., 2015; Miconi, 2016), and that this information may be represented in the orbitofrontal cortex (Wallis, 2007) or basal ganglia (Doya, 2000). Previous schemes, however, did not propose how this baseline “critic” may be instantiated, instead implementing it algorithmically. Here we propose a simple neural implementation of the baseline that automatically depends on the stimulus and thus does not require the learning system to have access to the true trial type, which in general is not known with certainty to the agent.

### Tasks with simple input-output mappings

The training procedure described in the previous section can be used for a variety of tasks, and results in networks that qualitatively reproduce both behavioral and electrophysiological findings from experiments with behaving animals. For the example perceptual decision-making task above, the trained network learns to integrate the sensory evidence to make the correct decision about which of two noisy inputs is larger (Figure 1C). This and additional networks trained for the same task were able to reach the target performance in ∼7,000 trials starting from completely random connection weights, and moreover the networks learned the “core” task after ∼2,000 trials (Figure 1—figure supplement 1). As with monkeys performing the task, longer stimulus durations allow the network to improve its performance by continuing to integrate the incoming sensory evidence (Wang, 2002; Kiani et al., 2008). Indeed, the output of the value network shows that the expected reward (in this case equivalent to confidence) is modulated by stimulus difficulty and increases in time (Figure 1—figure supplement 2); even though a confidence judgment is not strictly required for this task, the baseline value network provides a natural mechanism for extracting information about future reward from the policy network. This gives support to the hypothesis that confidence estimates are a fundamental and ubiquitous component of decision-making (Kepecs et al., 2008).

**Figure 2.**
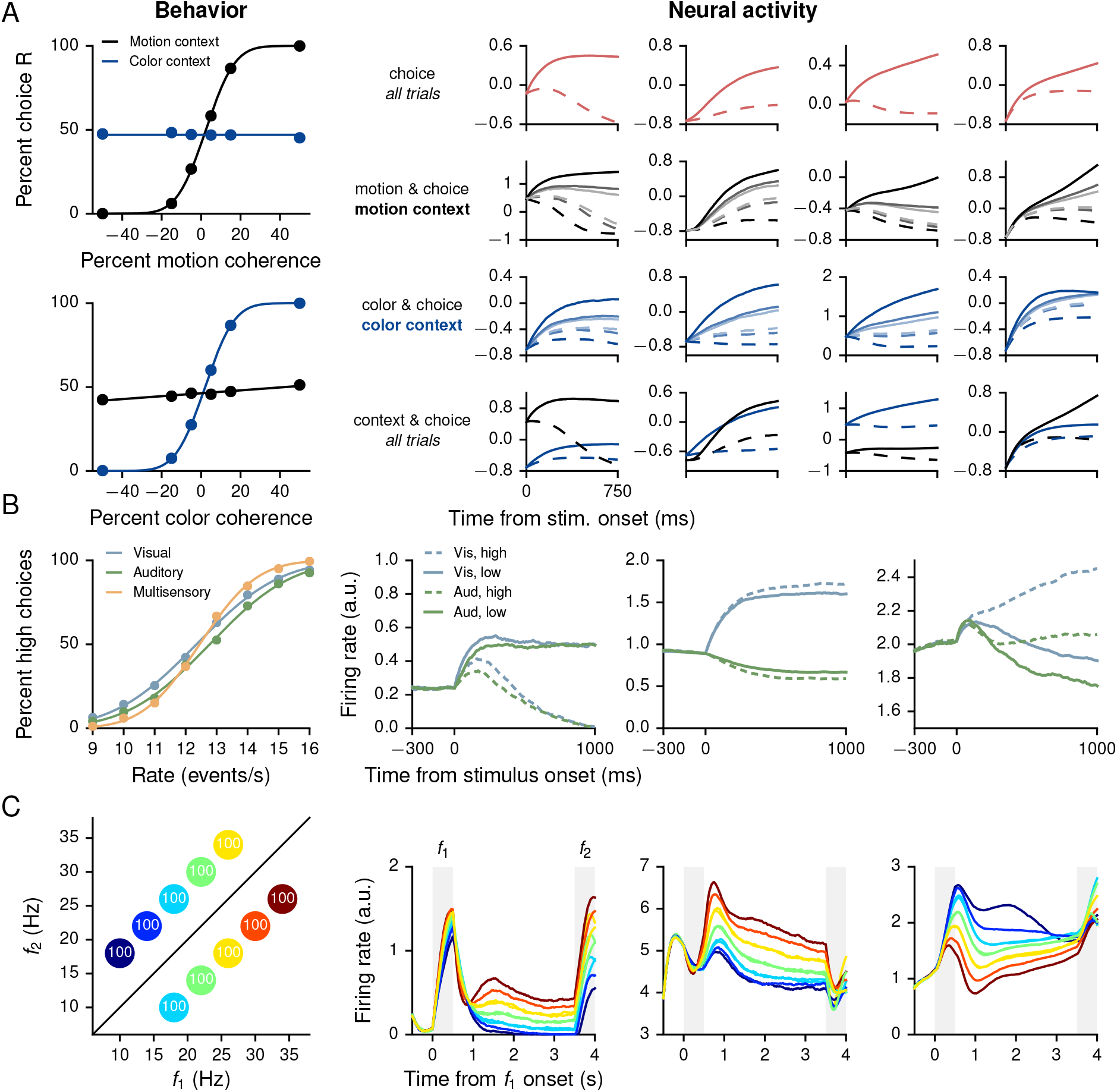
Performance and neural activity of RNNs trained for “simple” cognitive tasks in which the correct response depends only on the task condition. Left column shows behavioral performance, right column shows mixed selectivity for task parameters of example units in the policy network. (**A**) Contextdependent integration task (Mante et al., 2013). Left: Psychometric curves show the percentage of R choices as a function of the signed “motion” and “color” coherences. Right: Normalized firing rates of examples units sorted by different combinations of task parameters exhibit mixed selectivity. Firing rates were normalized by mean and standard deviation computed over the responses of all units, times, and trials. (B) Multisensory integration task (Raposo et al., 2012, 2014). Left: Psychometric curves show the percentage of high choices as a function of the event rate, for visual only (blue), auditory only (green), and multisensory (orange) trials. Improved performance on multisensory trials shows that the network learns to combine the two sources of information in accordance with Equation 13. Right: Sorted activity on visual only and auditory only trials for units selective for choice (high vs. low, left), modality (visual vs. auditory, middle), and both (right). (C) Parametric working memory task (Romo et al., 1999). Left: Percentage of correct responses for different combinations of *f*_1_ and *f*_2_. The conditions are colored here and in the right panels according to the first stimulus (base frequency) *f*_1_; due to the overlap in the values of *f*_1_, the 10 task conditions are represented by 7 distinct colors. Right: Activity of example policy network units sorted by *f*_1_. The first two units are positively tuned to *f*_1_ during the delay period, while the third unit is negatively tuned. Figure supplement 1. Learning curves for the context-dependent integration task (A). Figure supplement 2. Learning curves for the multisensory integration task (B). Figure supplement 3. Learning curves for the parametric working memory task (C).

Sorting the activity of individual units in the network by the signed coherence (the strength of the evidence, with negative values indicating evidence for L and positive for R) also reveals coherence-dependent ramping activity (Figure 1D) as observed in neural recordings from numerous perceptual decision-making experiments, e.g., Roitman & Shadlen (2002). The reaction time as a function of coherence in the reaction-time version of the same task, in which the go cue coincides with the time of stimulus onset, is also shown in Figure 1—figure supplement 3 and may be compared, e.g., to Wang (2002); Mazurek et al. (2003);Wong & Wang (2006). We note that, unlike in many neural models [e.g., Wong & Wang (2006)], here a firing-rate threshold is unnecessary for determining the reaction time; instead, the time at which the network commits to a decision is unambiguously given by the time at which the selected action is L or R. Learning curves for this and additional networks trained for the same reaction-time task are shown in Figure 1—figure supplement 4.

**Figure 3.**
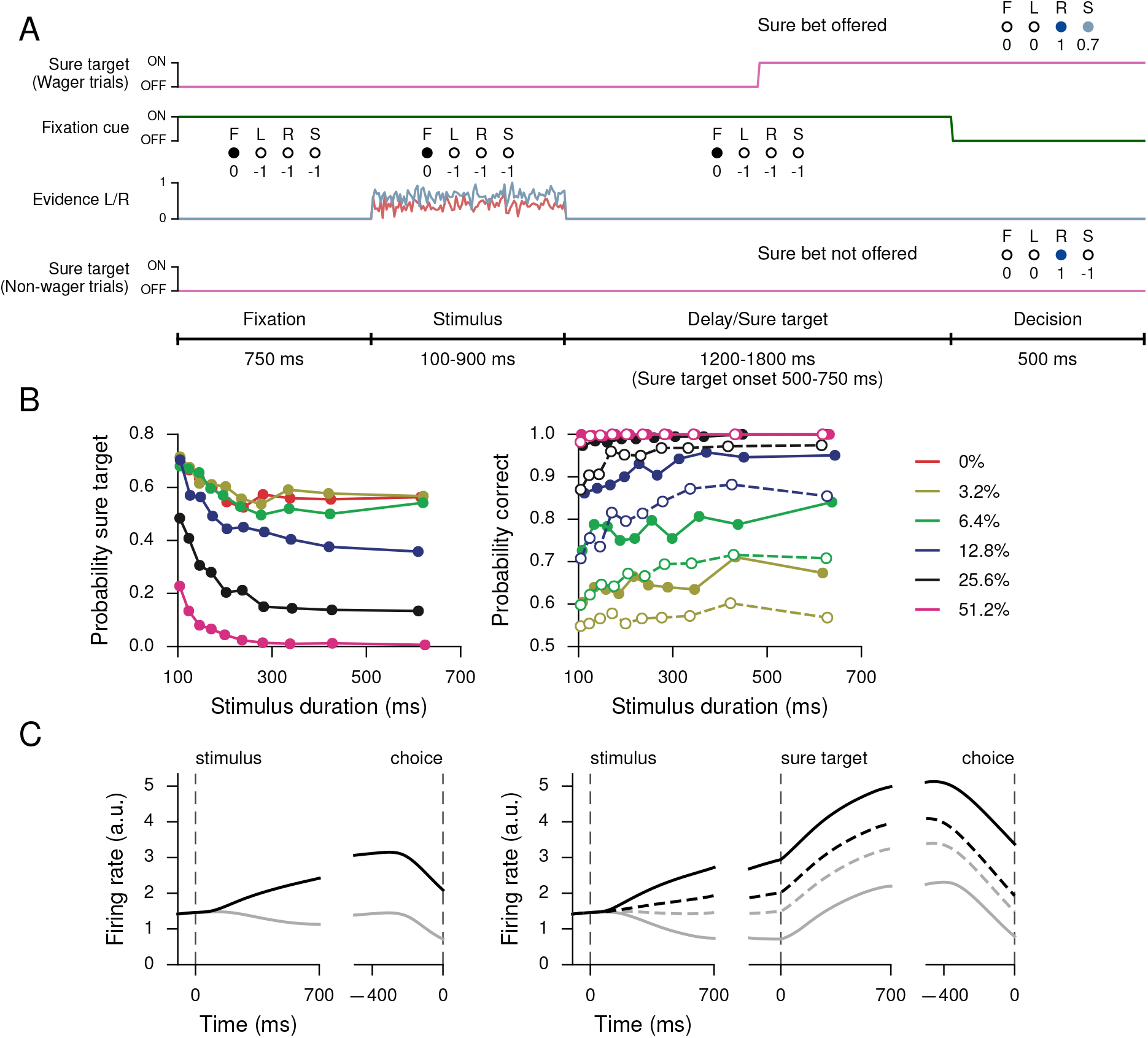
Perceptual decision-making task with postdecision wagering, based on Kiani & Shadlen (2009). (**A**) Task structure. On a random half of the trials, a sure option is presented during the delay period, and on these trials the network has the option of receiving a smaller (compared to correctly choosing L or R) but certain reward by choosing the sure option (S). The stimulus duration, delay, and sure target onset time are the same as in Kiani & Shadlen (2009). (**B**) Probability of choosing the sure option (left) and probability correct (right) as a function of stimulus duration, for different coherences. Performance is higher for trials on which the sure option was offered but waived in favor of L or R (filled circles, solid), compared to trials on which the sure option was not offered (open circles, dashed). (**C**) Activity of an example policy network unit for non-wager (left) and wager (right) trials, sorted by whether the presented evidence was toward the unit’s preferred (black) or nonpreferred (gray) target as determined by activity during the stimulus period on all trials. Dashed lines show activity for trials in which a sure option was chosen. Figure supplement 1. Learning curves.

**Figure 4.**
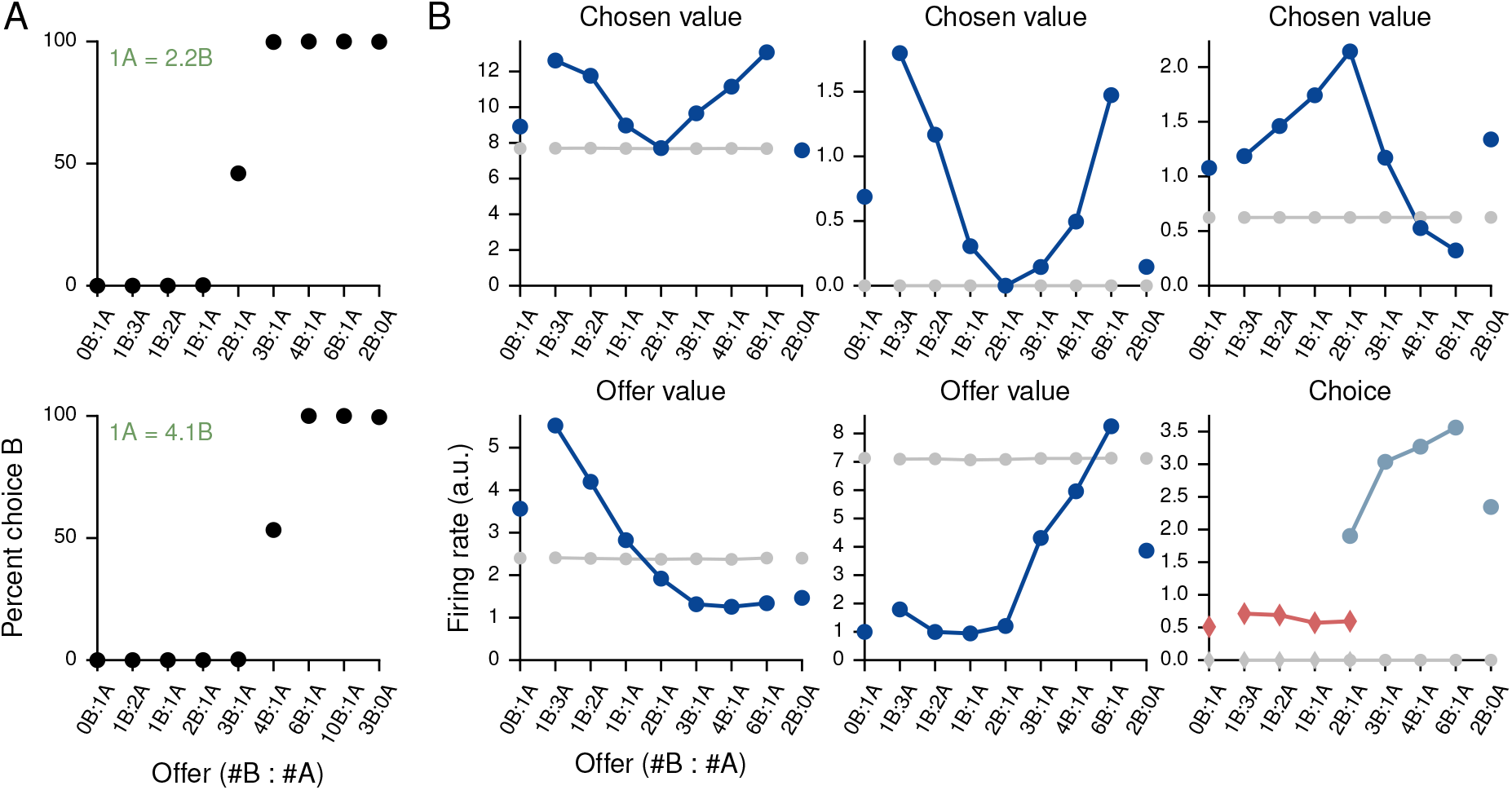
Value-based economic choice task (Padoa-Schioppa & Assad, 2006). (**A**) Choice pattern when the reward contingencies are indifferent for roughly 1 “juice” of A and 2 “juices” of B (upper) or 1 juice of A and 4 juices of B (lower). (**B**) Mean activity of example value network units during the pre-choice period, defined here as the period 500 ms before the decision, for the 1A = 2B case. Units in the value network exhibit diverse selectivity as observed in the monkey orbitofrontal cortex. For “choice” (last panel), trials were separated into choice A (red diamonds) and choice B (blue circles). Figure supplement 1. Fit of cumulative Gaussian to the choice pattern in (A), upper. Figure supplement 2. Fit of cumulative Gaussian to the choice pattern in (A), lower. Figure supplement 3. Learning curves.

In addition to the example task from the previous section, we trained networks for three well-known behavioral paradigms in which the correct, or optimal, behavior is determined on each trial by the task condition alone. Similar tasks have previously been addressed with several different forms of supervised learning, including FORCE (Sussillo & Abbott, 2009; Carnevale et al., 2015), Hessian-free (Martens & Sutskever, 2011; Mante et al., 2013; Barak et al., 2013), and stochastic gradient descent (Pascanu et al., 2013b; Song et al., 2016), so that the results shown in Figure 2 are presented as confirmation that the same tasks can also be learned using reward feedback on definite actions alone. For all three tasks the pre-stimulus fixation period was 750 ms; the networks had to maintain fixation until the start of a 500-ms “decision” period, which was indicated by the extinction of the fixation cue. At this time the network was required to choose one of two alternatives to indicate its decision and receive a reward of +1 for a correct response and 0 for an incorrect response; otherwise, the networks received a reward of -1.

The context-dependent integration task (Figure 2A) is based on Mante et al. (2013), in which monkeys were required to integrate one type of stimulus (the motion or color of the presented dots) while ignoring the other depending on a context cue. In training the network, we included both the 750-ms stimulus period and 300-1500-ms delay period following stimulus presentation. The delay consisted of 300 ms followed by a variable duration drawn from an exponential distribution with mean 300ms and truncated at amaximumof 1200 ms. The network successfully learned to perform the task, which is reflected in the psychometric functions showing the percentage of trials on which the network chose R as a function of the signed motion and color coherences, where motion and color indicate the two sources of noisy information and the sign is positive for R and negative for L (Figure 2A, left). As in electrophysiological recordings, units in the policy network show mixed selectivity when sorted by different combinations of task variables (Figure 2A, right). Learning curves for this and additional networks trained for the task are shown in Figure 2—figure supplement 1.

The multisensory integration task (Figure 2B) is based on Raposo et al. (2012, 2014), in which rats used visual flashes and auditory clicks to determine whether the event rate was higher or lower than a learned threshold of 12.5 events per second. When both modalities were presented, they were congruent, which implied that the rats could improve their performance by combining information from both sources. As in the experiment, the network was trained with a 1000-ms stimulus period, with inputs whose magnitudes were proportional (both positively and negatively) to the event rate. For this task the input connection weights *W*_in_, 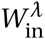*¸* and 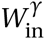 were initialized so that a third of the *N* = 150 policy network units received visual inputs only, another third auditory inputs only, and the remaining third received neither. As shown in the psychometric function (percentage of high choices as a function of event rate, Figure 2B, left), the trained network exhibits multisensory enhancement in which performance on multisensory trials was better than on single-modality trials. Indeed, like rats, the network combined the two modalities “optimally,” with

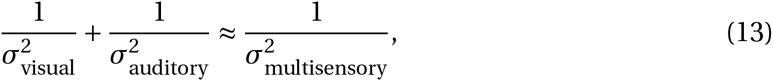

where 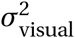, 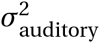, 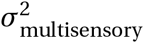 and are the variances obtained from fits of the psychometric functions to cumulative Gaussian functions for visual only, auditory only, and multisensory (both visual and auditory) trials, respectively (Table 2). As observed in electrophysiological recordings, moreover, policy network units exhibit a wide range of tuning to task parameters, with some selective to choice and others to modality, while mawny units showed mixed selectivity to all task variables (Figure 2B, right). Learning curves for this and additional networks trained for the task are shown in Figure 2—figure supplement 2.

**Table 2.** Psychophysical thresholds *σ*_visual_, *σ*_auditory_, and *σ*_multisensory_ obtained from fits of cumulative Gaussian functions to the psychometric curves in visual only, auditory only, and multisensory trials in the multisensory integration task, for 6 networks trained from different random initializations (first row, bold: network from main text, cf. Figure 2B). The last two columns show evidence of “optimal” multisensory integration according to Equation 13 (Raposo et al., 2012).

| $\sigma_{\text{visual}}$ | $\sigma_{\text{auditory}}$ | $\sigma_{\text{multisensory}}$ | $\frac{1}{\sigma_{\text{visual}}^2} + \frac{1}{\sigma_{\text{auditory}}^2}$ | $\frac{1}{\sigma_{\text{multisensory}}^2}$ |
| --- | --- | --- | --- | --- |
| **2.124** | **2.099** | **1.451** | **0.449** | **0.475** |
| 2.107 | 2.086 | 1.448 | 0.455 | 0.477 |
| 2.276 | 2.128 | 1.552 | 0.414 | 0.415 |
| 2.118 | 2.155 | 1.508 | 0.438 | 0.440 |
| 2.077 | 2.171 | 1.582 | 0.444 | 0.400 |
| 2.088 | 2.149 | 1.480 | 0.446 | 0.457 |

The parametric working memory task (Figure 2C) is based on the vibrotactile frequency discrimination task of Romo et al. (1999), in which monkeys were required to compare the frequencies of two temporally separated stimuli to determine which was higher. For network training, the task epochs consisted of a 500-ms base stimulus with “frequency” *f*_1_, a 2700-3300-ms delay, and a 500-ms comparison stimulus with frequency *f*_2_; for the trials shown in Figure 2C the delay was always 3000 ms as in the experiment. During the decision period, the network had to indicate which stimulus was higher by choosing *f*_1_ < *f*_2_ or *f*_1_ > *f*_2_. The stimuli were constant inputs with amplitudes proportional (both positively and negatively) to the frequency. For this task we set the learning rate to *η* = 0.002; the network successfully learned to perform the task (Figure 2C, left), and the individual units of the network, when sorted by the first stimulus (base frequency) *f*_1_, exhibit highly heterogeneous activity (Figure 2C, right) characteristic of neurons recorded in the prefrontal cortex of monkeys performing the task (Machens et al., 2010). Learning curves for this and additional networks trained for the task are shown in Figure 2—figure supplement 3.

Additional comparisons can be made between the model networks shown in Figure 2 and the neural activity observed in behaving animals, for example state-space analyses as in Mante et al. (2013), Carnevale et al. (2015), or Song et al. (2016). Such comparisons reveal that, as found previously in studies such as Barak et al. (2013), the model networks exhibit many, but not all, features present in electrophysiological recordings. Figure 2 and the following make clear, however, that RNNs trained with reward feedback alone can already reproduce the mixed selectivity characteristic of neural populations in higher cortical areas (Rigotti et al., 2010, 2013), thereby providing a valuable platform for future investigations of how such complex representations are learned.

### Confidence and perceptual decision-making

All of the tasks in the previous section have the property that the correct response on any single trial is a function only of the task condition, and, in particular, does not depend on the network’s state during the trial. In a postdecision wager task (Kiani & Shadlen, 2009), however, the optimal decision depends on the animal’s (agent’s) estimate of the probability that its decision is correct, i.e., its confidence. This makes it difficult to train with standard supervised learning: instead, we trained an RNN to perform the task by maximizing overall reward. On a random half of the trials a “sure” option was presented, and selecting this option resulted in a reward that was 0.7 times the size of the reward obtained when correctly choosing L or R. As in the monkey experiment, the network receives no information indicating whether or not a given trial will contain a sure option until the middle of the delay period after stimulus offset, thus ensuring that the network makes a decision about the stimulus on all trials (Figure 3A). For this task the input connection weights *W*_in_, 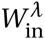*¸* and 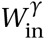 were initialized so that half the units received information about the sure target while the other half received evidence for L and R. All units initially received fixation input.

The key behavioral features found in Kiani & Shadlen (2009); Wei & Wang (2015) are reproduced in the trained network, namely the network opted for the sure option more frequently when the coherence was small or stimulus duration short (Figure 3B, left); and when the network was presented with a sure option but waived it in favor of choosing L or R, the performance was better than on trials when the sure option was not presented (Figure 3B, right). The latter observation is taken as indication that neither monkeys nor trained networks choose the sure target on the basis of stimulus difficulty alone but based on their internal sense of uncertainty on each trial.

Figure 3C shows the activity of an example network unit, sorted by whether the decision was the unit’s preferred or nonpreferred target (as determined by firing rates during the stimulus period on all trials), for both non-wager and wager trials. In particular, on trials in which the sure option was chosen, the firing rate is intermediate compared to trials on which the network made a decision by choosing L or R. Learning curves for this and additional networks trained for the task are shown in Figure 3—figure supplement 1.

### Value-based economic choice task

We also trained networks to perform the simple economic choice task of Padoa-Schioppa & Assad (2006) and examined the activity of the *value*, rather than policy, network. The choice patterns of the networks were modulated only by varying the reward contingencies (Figure 4A, upper and lower). We note that, on each trial there is a “correct” answer in the sense that there is a choice which results in greater reward. In contrast to the previous tasks, however, information regarding whether an answer is correct in this sense is not contained in the inputs but rather in the association between inputs and rewards. This distinguishes the task from the cognitive tasks discussed in previous sections.

Each trial began with a 750-ms fixation period; the offer, which indicated the “juice” type and amount for the left and right choices, was presented for 1000-2000 ms, followed by a 750-ms decision period during which the network was required to indicate its decision. In the upper panel of Figure 4A the indifference point was set to 1A = 2.2B during training, which resulted in 1A = 2.0B when fit to a cumulative Gaussian (Figure 4—figure supplement 1), while in the lower panel it was set to 1A = 4.1B during training and resulted in 1A = 4.0B (Figure 4—figure supplement 2). The basic unit of reward, i.e., 1B, was 0.1. For this task we increased the initial value of the value network’s input weights, 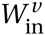, by a factor of 10 to drive the value network more strongly.

Strikingly, the activity of units in the value network 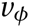 exhibits similar types of tuning to task variables as observed in the orbitofrontal cortex of monkeys, with some units (roughly 20% of active units) selective to chosen value, others (roughly 60%, for both A and B) to offer value, and still others (roughly 20%) to choice alone as defined in Padoa-Schioppa & Assad (2006) (Figure 4B). The policy network also contained units with a diversity of tuning. Learning curves for this and additional networks trained for the task are shown in Figure 4—figure supplement 3. We emphasize that no changes were made to the network architecture for this value-based economic choice task. Instead, the same scheme shown in Figure 1B, in which the value network is responsible for predicting future rewards to guide learning but is *not* involved in the execution of the policy, gave rise to the pattern of neural activity shown in Figure 4B.

## Discussion

In this work we have demonstrated reward-based training of recurrent neural networks for both cognitive and value-based tasks. Our main contributions are twofold: first, our work expands the range of tasks and corresponding neural mechanisms that can be studied by analyzing model recurrent neural networks, while providing a unified setting in which to study such diverse computations and compare to electrophysiological recordings from behaving animals; second, by explicitly incorporating reward into network training, our work makes it possible to more directly address the question of reward-dependent processes in the brain, for instance the role of value representation that is essential for learning, but not for executing, a task.

To our knowledge, the specific form of the baseline network inputs used in this work, which was chosen as a biologically plausible candidate architecture for implementing policy search reinforcement learning, has not been used previously in the context of recurrent policy gradients; it combines ideas from Wierstra et al. (2009) where the baseline network received the same inputs as the policy network in addition to the selected actions, and Ranzato et al. (2016), where the baseline was implemented as a simple linear regressor of the activity of the policy network so that the policy and value networks effectively shared the same recurrent units. Indeed, for some of the simpler tasks considered here, models with a baseline consisting of a linear readout of the selected actions and policy network activity could be trained in comparable (but slightly longer) time (Figure 1—figure supplement 5). A separate baseline network, however, suggests an explicit role for value representation in the brain that is essential for learning a task (equivalently, if the environment is changing), but not for executing an already learned task, as is sometimes found in experiments (Turner & Desmurget, 2010; Schoenbaum et al., 2011; Stalnaker et al., 2015). Since an accurate baseline dramatically improves learning but is not *required*—the algorithm takes many more samples to converge with a constant baseline, for instance—this baseline network hypothesis for the role of value representation may account for some of the subtle yet broad learning deficits observed in OFC-lesioned animals (Wallis, 2007). Moreover, since expected reward is closely related to decision confidence in many of the tasks considered, a value network that nonlinearly reads out confidence information from the policy network is consistent with experimental findings in which OFC inactivation affects the ability to report confidence but not decision accuracy (Lak et al., 2014).

Our results thus support the actor-critic picture for reward-based learning, in which one circuit directly computes the policy to be followed, while a second structure, receiving projections from the policy network as well as information about the selected actions, computes expected future reward to guide learning. We note that this is the opposite of the way value functions are usually thought of in neuroscience, which stresses the primacy of the value function from which policies are derived. However, there is experimental evidence that signals in the striatum are more suitable for direct policy search rather than for updating action values as an intermediate step, as would be the case for value function-based approaches to computing the policy (Li & Daw, 2011; Niv & Langdon, 2016). Moreover, although we have used a single RNN each to represent the policy and value modules, using “deep,” multilayer RNNs may increase the representational power of each module (Pascanu et al., 2013a). For instance, more complex tasks than considered in this work may require hierarchical feature representation in the policy network, and likewise value networks can use a combination of the different features [including raw sensory inputs (Wierstra et al., 2009)] to predict future reward. Anatomically, the policy networks may correspond to circuits in dorsolateral prefrontal cortex, while the value networks may correspond to circuits in the orbitofrontal cortex (Schultz et al., 2000; Takahashi et al., 2011) or basal ganglia (Hikosaka et al., 2014). This architecture also provides a useful example of the hypothesis that various areas of the brain effectively optimize different cost functions (Marblestone et al., 2016): in this case, the policy network maximizes reward, while the value network minimizes the prediction error for future reward.

As in many other supervised learning approaches used previously to train RNNs (Mante et al., 2013; Song et al., 2016), the use of BPTT to compute the gradients (in particular, the eligibility) make our “plasticity rule” not biologically plausible. Thus our focus has been on learning from realistic feedback signals provided by the environment but not on its physiological implementation. Nevertheless, recent work suggests that exact backpropagation is not necessary (Lillicrap et al., 2014), and approximate forms of backpropagation and SGD can be implemented in a biologically plausible manner (Scellier & Bengio, 2016). Such ideas require further investigation and may lead to effective yet more neurally plausible methods for training model neural networks.

Recently, Miconi (2016) used a “node perturbation”-based (Fiete & Seung, 2006; Fiete et al., 2007; Hoerzer et al., 2014) algorithm with an error signal at the end of each trial to train RNNs for several cognitive tasks, and indeed, node perturbation is closely related to the REINFORCE algorithm used in this work. On one hand, the method described in Miconi (2016) is more biologically plausible in the sense of not requiring gradients computed via backpropagation through time as in our approach; on the other hand, in contrast to the networks in this work, those in Miconi (2016) did not “commit” to a discrete action and thus the error signal was a graded quantity. In this and other works (Frémaux et al., 2010), moreover, the prediction error was computed by algorithmically keeping track of a stimulus (task condition)-specific running average of rewards. Here we propose a simple, concrete scheme for approximating the average that automatically depends on the stimulus, without requiring an external learning system to maintain a separate record for each (true) trial type, which is not known by the agent with certainty.

One of the advantages of the REINFORCE algorithm for policy gradient reinforcement learning is that direct supervised learning can also be mixed with reward-based learning, by including only the eligibility term in Equation 3 without modulating by reward (Mnih et al., 2014), i.e., by maximizing the log-likelihood of the desired actions. Although all of the networks in this work were trained from reward feedback only, it will be interesting to investigate this feature of the REINFORCE algorithm in the context of human learning. Another advantage, which we have not exploited here, is the possibility of learning policies for continuous action spaces (Peters & Schaal, 2008; Wierstra et al., 2009); this would allow us, for example, to model arbitrary saccade targets in the perceptual decision-making task, rather than limiting the network to discrete choices.

We have previously emphasized the importance of incorporating biological constraints in the training of neural networks (Song et al., 2016). For instance, neurons in the mammalian cortex have purely excitatory or inhibitory effects on other neurons, which is a consequence of Dale’s Principle for neurotransmitters (Eccles et al., 1954). In this work we did not include such constraints due to the more complex nature of our rectified GRUs (Equations 9-12); in particular, the units we used are capable of dynamically modulating their time constants and gating their recurrent inputs, and we therefore interpreted the firing rate units as a mixture of both excitatory and inhibitory populations. Indeed, these may implement the “reservoir of time constants” observed experimentally (Bernacchia et al., 2011). In the future, however, comparison to both model spiking networks and electrophysiological recordings will be facilitated by including more biological realism, by explicitly separating the roles of excitatory and inhibitory units (Mastrogiuseppe & Ostojic, 2016). Moreover, since both the policy and value networks are obtained by minimizing an objective function, additional regularization terms can be easily included to obtain networks whose activity is more similar to neural recordings (Sussillo et al., 2015; Song et al., 2016).

Finally, one of the most appealing features of RNNs trained to perform many tasks is their ability to provide insights into neural computation in the brain. However, methods for revealing neural mechanisms in such networks remain limited to state-space analysis (Sussillo & Barak, 2013), which in particular does not reveal how the synaptic connectivity leads to the dynamics responsible for implementing the higher-level policy. General and systematic methods for analyzing trained networks are still needed and are the subject of ongoing investigation. Nevertheless, reward-based training of RNNs makes it more likely that the resulting networks will correspond closely to biological networks observed in experiments with behaving animals. We expect that the continuing development of tools for training model neural networks in neuroscience will thus contribute novel insights into the neural basis of animal cognition.

## Methods

### Policy gradient reinforcement learning with RNNs

Here we review the application of the REINFORCE algorithm for policy gradient reinforcement learning to recurrent neural networks (Williams, 1992; Sutton & Barto, 1998; Peters & Schaal, 2008; Wierstra et al., 2009). In particular, we provide a careful derivation of Equation 3 following, in part, the exposition in Zaremba & Sutskever (2016).

Let *H*_*μ*:*t*_ be the sequence of interactions between the environment and agent (i.e., the environmental states, observables, and agent actions) that results in the environment being in state **s**_*t*+1_ at time *t* +1 starting from state **s**_*μ*_ at time *μ*:

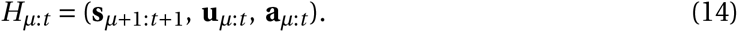

For notational convenience in the following, we adopt the convention that, for the special case of *μ* = 0, the history *H*_0:*t*_ includes the initial state **s**_0_ and excludes the meaningless inputs **u**_0_, which are not seen by the agent:

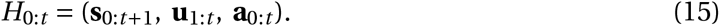

When *t* = 0, it is also understood that 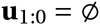, the empty set. A full history, or a trial, is thus denoted as

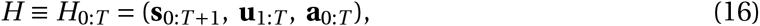

where *T* is the end of the trial. Indeed, since different trials can have different durations, we take *T* to be the maximum length of a trial in the task. The reward *ρ*_*t*+1_ at time *t* +1 following actions **a**_*t*_ (we use *ρ* to distinguish it from the firing rates **r** of the RNNs) is determined by this history, which we sometimes indicate explicitly by writing

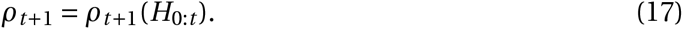

As noted in the main text, we adopt the convention that the agent performs actions at *t* = 0,…,*T* and receives rewards at *t* = 1,…,*T* +1 to emphasize that rewards follow the actions and are jointly determined with the next state (Sutton& Barto, 1998). For notational simplicity, here and elsewhere we assume that any discount factor is already included in *ρ*_*t*+1_, i.e., in all places where the reward appears we consider 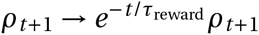, where *τ*_reward_ is the time constant for discounting future rewards (Doya, 2000); we included temporal discounting only for the reaction-time version of the simple perceptual decision-making task (Figure 1—figure supplement 3), where we set *τ*_reward_ = 10 s. For the remaining tasks, 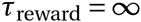.

Explicitly, a trial *H*_0:*T*_ comprises the following. At time *t* = 0, the environment is in state **s**_0_ with probability ℰ (**s**_0_). The agent initially chooses a set of actions **a**_0_ with probability 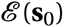, which is determined by the parameters of the policy network, in particular the initial conditions **x**_0_ and readout weights 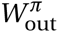 and biases 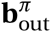 (Equation 6). At time *t* = 1, the environment, depending on its previous state **s**_0_ and the agent’s actions **a**_0_, transitions to state **s**_1_ with probability ℰ (**s**_1_|**s**_0_,**a**_0_). The history up to this point is 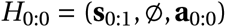, where 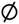 indicates that no inputs have yet been seen by the network. The environment also generates reward *ρ*_1_, which depends on this history, 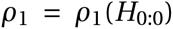. From state **s**_1_ the environment generates observables (inputs to the agent) **u**_1_ with a distribution given by 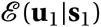. In response, the agent, depending on the inputs **u**_1_ it receives from the environment, chooses the set of actions **a**_1_ according to the distribution 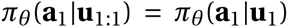. The environment, depending on its previous states **s**_0:1_ and the agent’s previous actions **a**_0:1_, then transitions to state **s**_2_ with probability 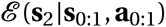. Thus 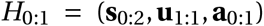. Iterating these steps, the history at time *t* is therefore given by Equation 15, while a full history is given by Equation 16.

The probability 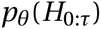 of a particular sub-history 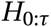 up to time *τ* occurring, under the policy *π*_*θ*_ parametrized by *θ*, is given by

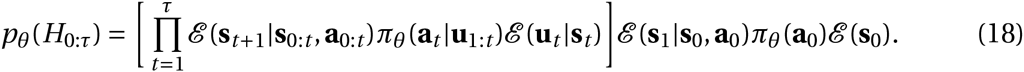

In particular, the probability *p*_*θ*_(*H*) of a history *H* = *H*_0:*T*_ occurring is

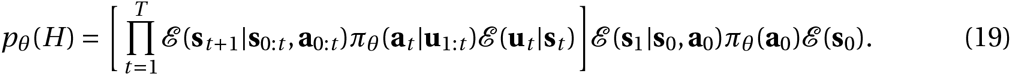

A key ingredient of the REINFORCE algorithm is that the logarithmic derivatives of Equation 18 with respect to the parameters *θ* do not depend on the unknown (to the agent) environmental dynamics contained in ℰ, because

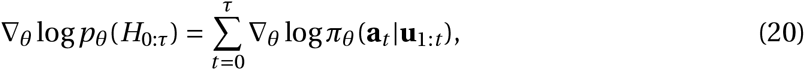

with the understanding that **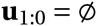** (the empty set) and therefore **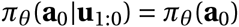**.

The goal of the agent is to maximize the expected return at time *t* = 0 (Equation 1, reproduced here)

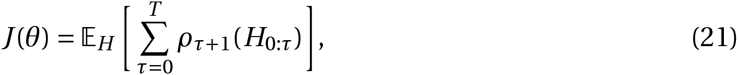

where we have used the time index *τ* for notational consistency with the following and made the history-dependence of the rewards explicit. In terms of the probability of each history *H* occurring, Equation 19, we have

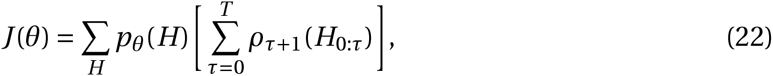

where the generic sum over *H* may include both sums over discrete variables and integrals over continuous variables. Using the “likelihood trick”

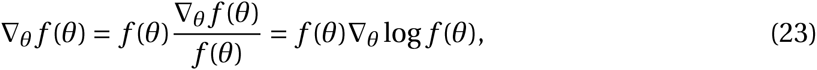

we can write

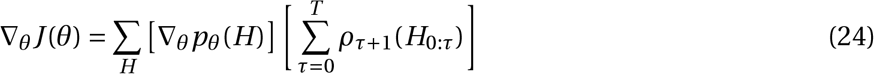

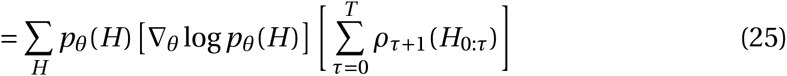

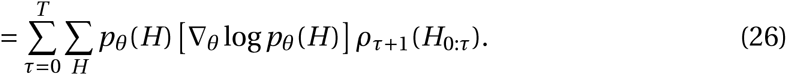

Since, for any *τ* = 0,…,*T*,

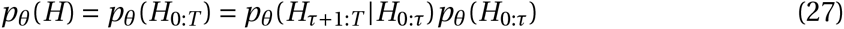

(cf. Equation 18), we can take the logarithm and insert into Equation 26 to obtain

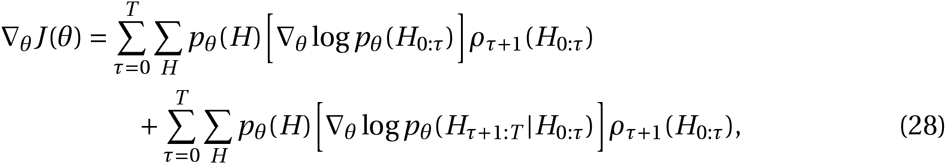

where the second termis zero because (again using Equation 27)

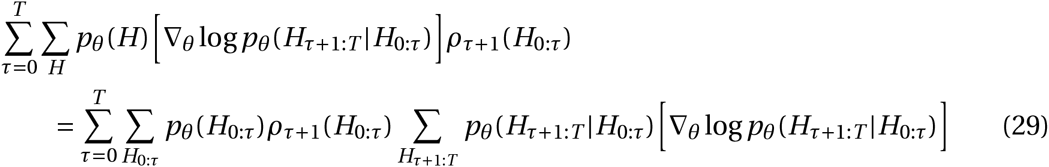

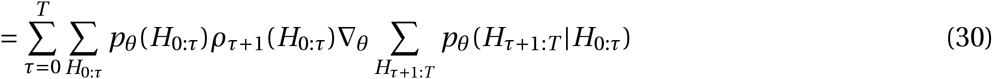

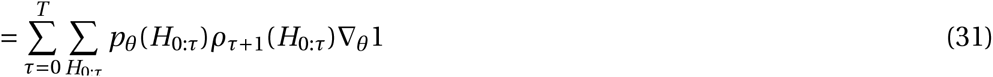

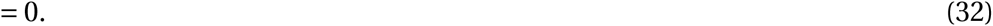

Therefore, from Equation 20 and the remaining termin Equation 28, we obtain

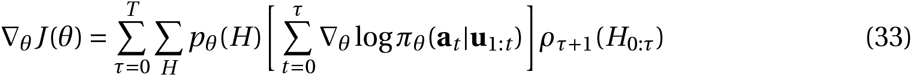

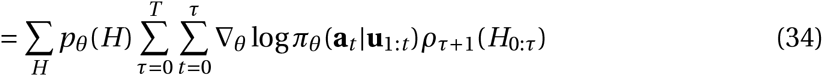

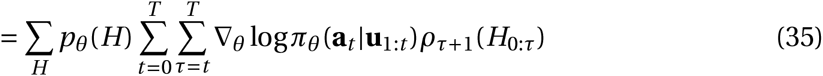

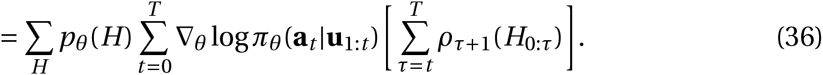

We then obtain the first terms of Equation 2 and Equation 3 by estimating the sum over all *H* by *N*_trials_ samples from the agent’s experience.

In Equation 22 it is evident that subtracting a constant baseline for each parameter will not affect the *gradient* of *J*(*θ*), i.e., Equation 36. It is possible to use this invariance to find an “optimal” value of the constant baseline that minimizes the variance of the gradient estimate (Peters & Schaal, 2008). In practice, however, it is more useful to have a history-dependent baseline that attempts to predict the future return at every time (Wierstra et al., 2009; Zaremba & Sutskever, 2016). We therefore introduce a second network, called the *value network*, that uses the selected actions **a**_1:*t*_ and the activity of the policy network 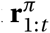 to predict the future return 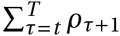 by minimizing the squared error (Equations 4-5). Intuitively, such a baseline is appealing because the terms in the gradient of Equation 3 are nonzero only if the actual return deviates from what was predicted by the value network.

### Discretized network equations and initialization

Carrying out the discretization of Equations 9-12 in time steps of Δ*t*, we obtain

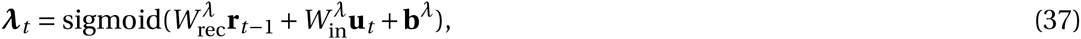

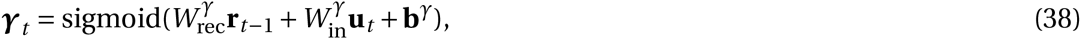

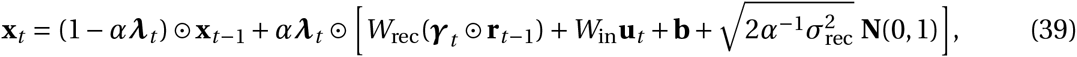

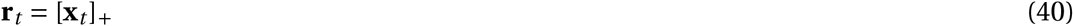

for *t* = 1,…,*T*, where *α* = Δ *t*/*τ* and **N**(0,1) are normally distributed random numbers with zero mean and unit variance.

The biases **b^λ^**, **b**^*γ*^, and **b**, as well as the readout weights 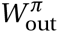 and 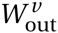, were initialized to zero. The biases for the policy readout 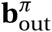 were initially set to zero, while the value network bias 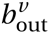 was initially set to the “reward” for an aborted trial, -1. The entries of the inputweight matrices 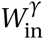, 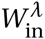*¸* and 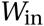 for both policy and value networks were drawn from a zero-mean Gaussian distribution with variance 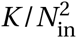. For the recurrent weight matrices *W*_rec_, 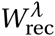*¸* and 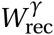, the *K* nonzero entries in each row were initialized from a gamma distribution Γ(*α, β*) with *α* = *β* = 4, with each entry multiplied randomly by ±1; the entire matrix was then scaled such that the spectral radius—the largest absolute value of the eigenvalues—was exactly *ρ*_0_. Although we also successfully trained networks starting from normally distributed weights, we found it convenient to control the sign and magnitude of the weights independently. The initial conditions **x**_0_, which are also trained, were set to 0.5 for all units before the start of training. We implemented the networks in the Python machine learning library Theano (The Theano Development Team, 2016).

### AdamSGD with gradient clipping

We use a recently developed version of stochastic gradient descent known as Adam, for *ada*ptive *m*oment estimation (Kingma & Ba, 2015), together with gradient clipping to prevent exploding gradients (Graves, 2013; Pascanu et al., 2013b). For clarity, in this section we use vector notation ***θ*** to indicate the set of all parameters being optimized and the subscript *k* to indicate a specific parameter *θ*_*k*_. At each iteration *i* > 0, let

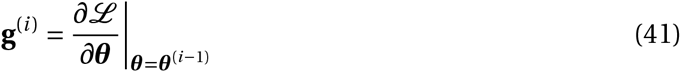

be the gradient of the objective function ℒ with respect to the parameters ***θ***. We first clip the gradient if its norm |**g**^(*i*)^| exceeds amaximum Γ (see Table 1), i.e.,

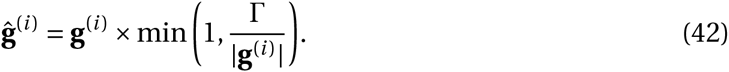

Each parameter *θ*_*k*_ is then updated according to

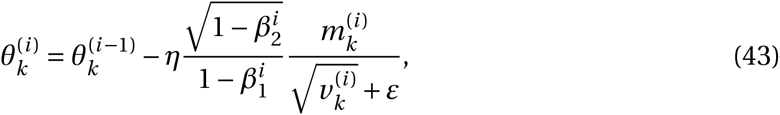

where *η* is the base learning rate and the moving averages

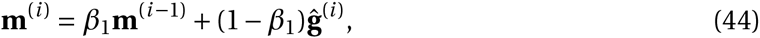

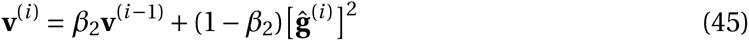

estimate the first and second (uncentered) moments of the gradient. Initially, **m**^(0)^ = **v**^(0)^ = 0. These moments allow each parameter to be updated in Equation 43 according to adaptive learning rates, such that parameters whose gradients exhibit high uncertainty and hence large “signal-to-noise ratio” lead to smaller learning rates.

Except for the base learning rate *η* (see Table 1), we used the parameter values suggested in Kingma & Ba (2015):

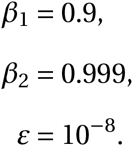

### Computer code

All code used in this work, including code for generating the figures, is available at http://github.com/xjwanglab/pyrl.

## Acknowledgements

We thank R. Pascanu, K. Cho, E. Ohran, and E. Russek for valuable discussions. This work was supported by Office of Naval Research Grant N00014-13-1-0297 and a Google Computational Neuroscience Grant.

## Author contributions

HFS, participated in all aspects of the project; GRY, analysis and interpretation of data, drafting and revising the article; X-JW, conception and design, analysis and interpretation of data, drafting and revising the article.

---

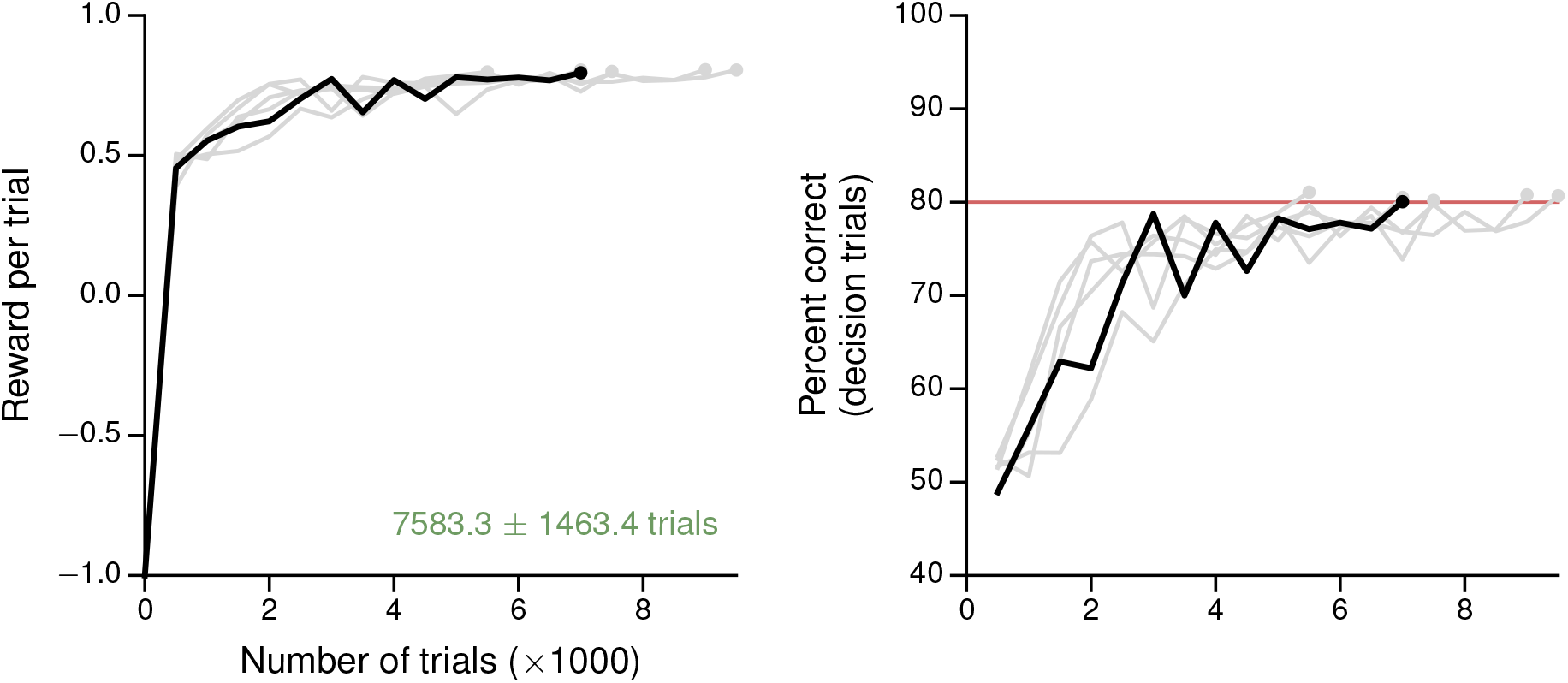
Figure 1—figure supplement 1. Learning curves for the simple perceptual decision-making task. (**A**) Average reward per trial. Black indicates the network realization shown in the main text, gray additional realizations, i.e., trained with different random number generator seeds. (**B**) Percent correct, for trials on which the network made a decision (≥ 99% required for termination). Red: target performance (when training was terminated).

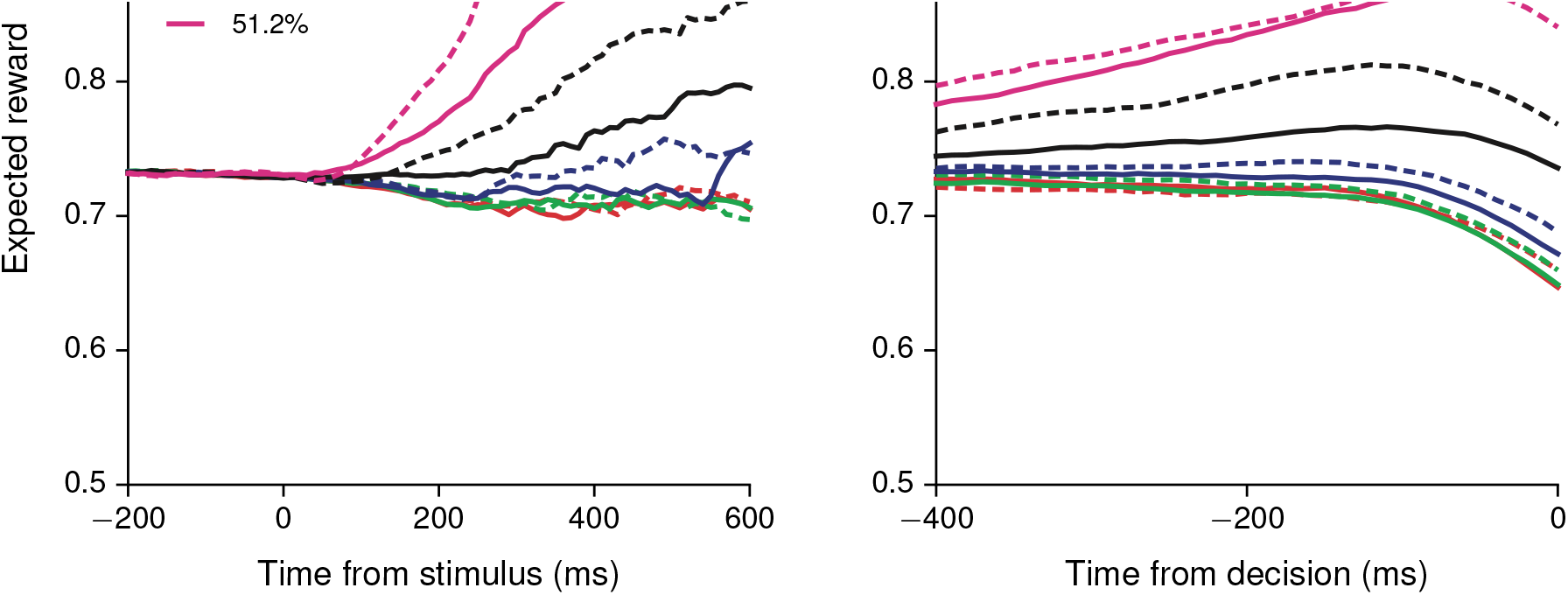
Figure 1—figure supplement 2. Output of the value network (expected reward) aligned to (**A**) stimulus onset and (**B**) choice, sorted by coherence for positive (solid) and negative (dashed) coherences.

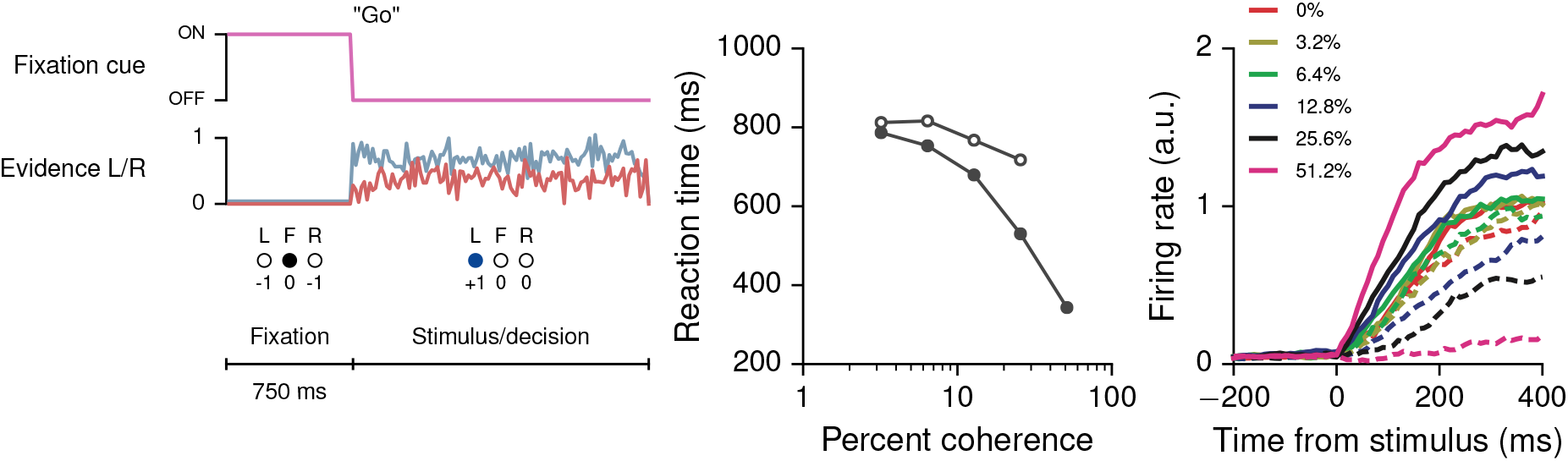
Figure 1—figure supplement 3. (**A**) Task structure for the reaction-time version of the simple perceptual decision-making task, in which the agent can choose to respond any time after the onset of stimulus. (**B**) Reaction time as a function of coherence for correct (solid circles) and error (open circles) trials. (**C**) Neural activity of an example policy network unit, sorted by the coherence (the difference in strength of evidence for L and R) and aligned to the time of stimulus onset. Each trial ends when the network breaks fixation.

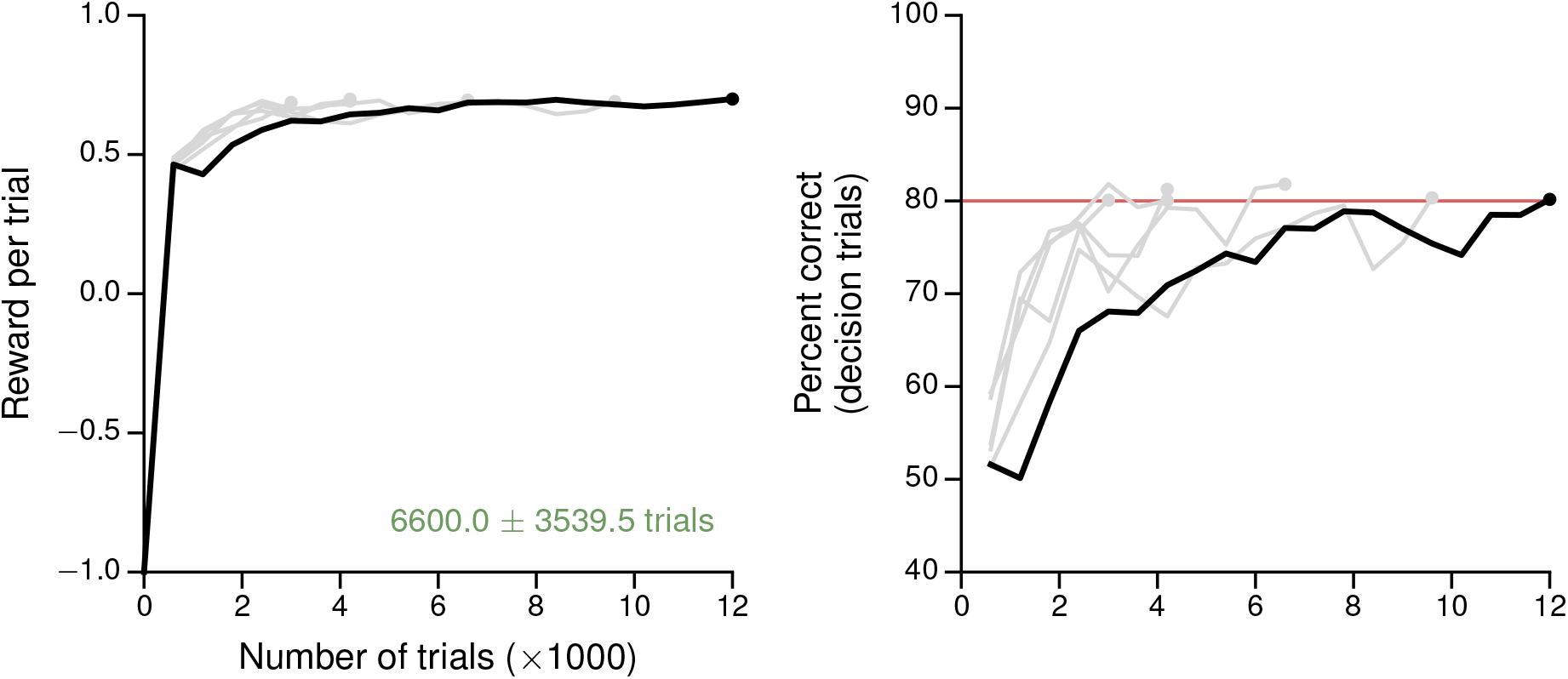
Figure 1—figure supplement 4. Learning curves for the reaction-time version of the simple perceptual decision-making task. (**A**) Average reward per trial. Black indicates the network realization shown in the main text, gray additional realizations, i.e., trained with different random number generator seeds. (**B**) Percent correct, for trials on which the network made a decision (≥ 99% required for termination). Red: target performance (when training was terminated).

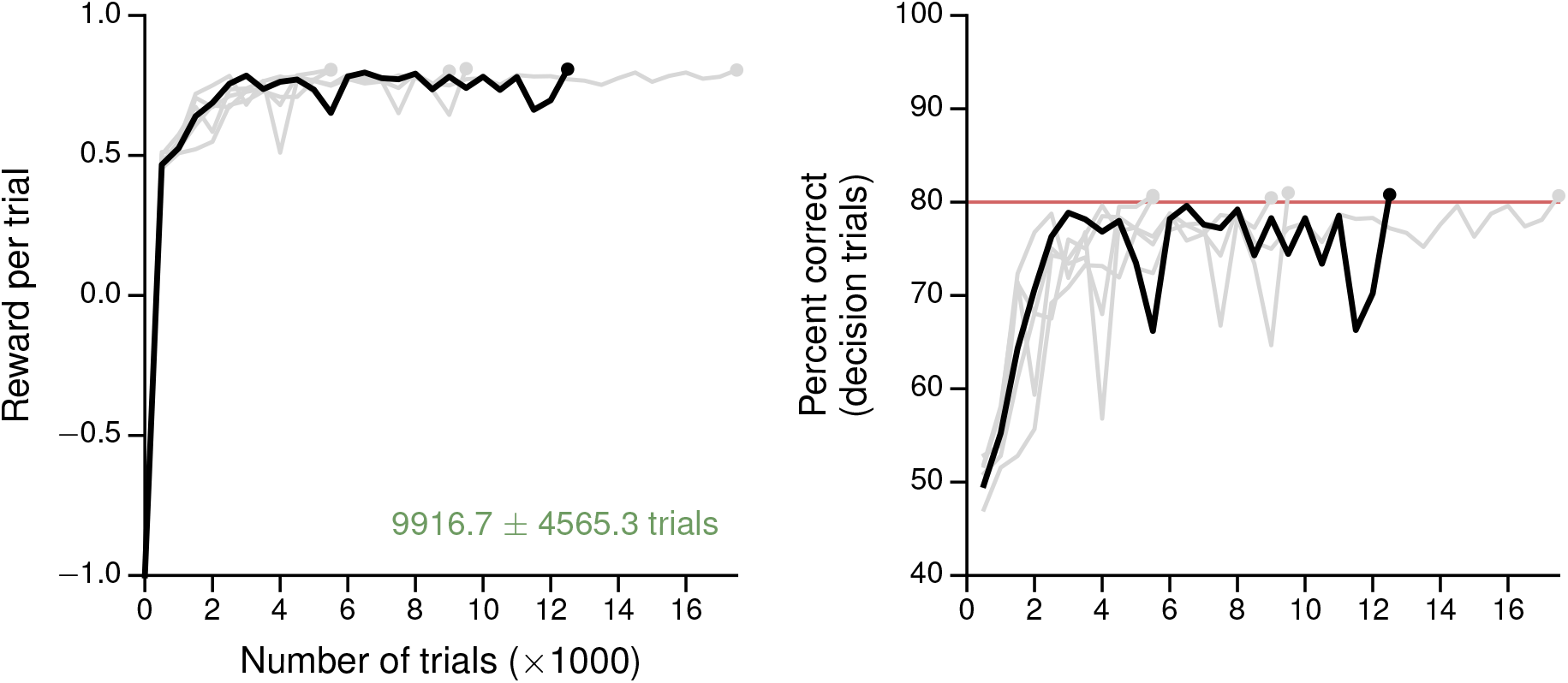
Figure 1—figure supplement 5. Learning curves for the simple perceptual decision-making task with a linear readout of the policy network as the baseline. (**A**) Average reward per trial. Black indicates the network realization shown in the main text, gray additional realizations, i.e., trained with different random number generator seeds. (**B**) Percent correct, for trials on which the network made a decision (≥ 99% required for termination). Red: target performance (when training was terminated).

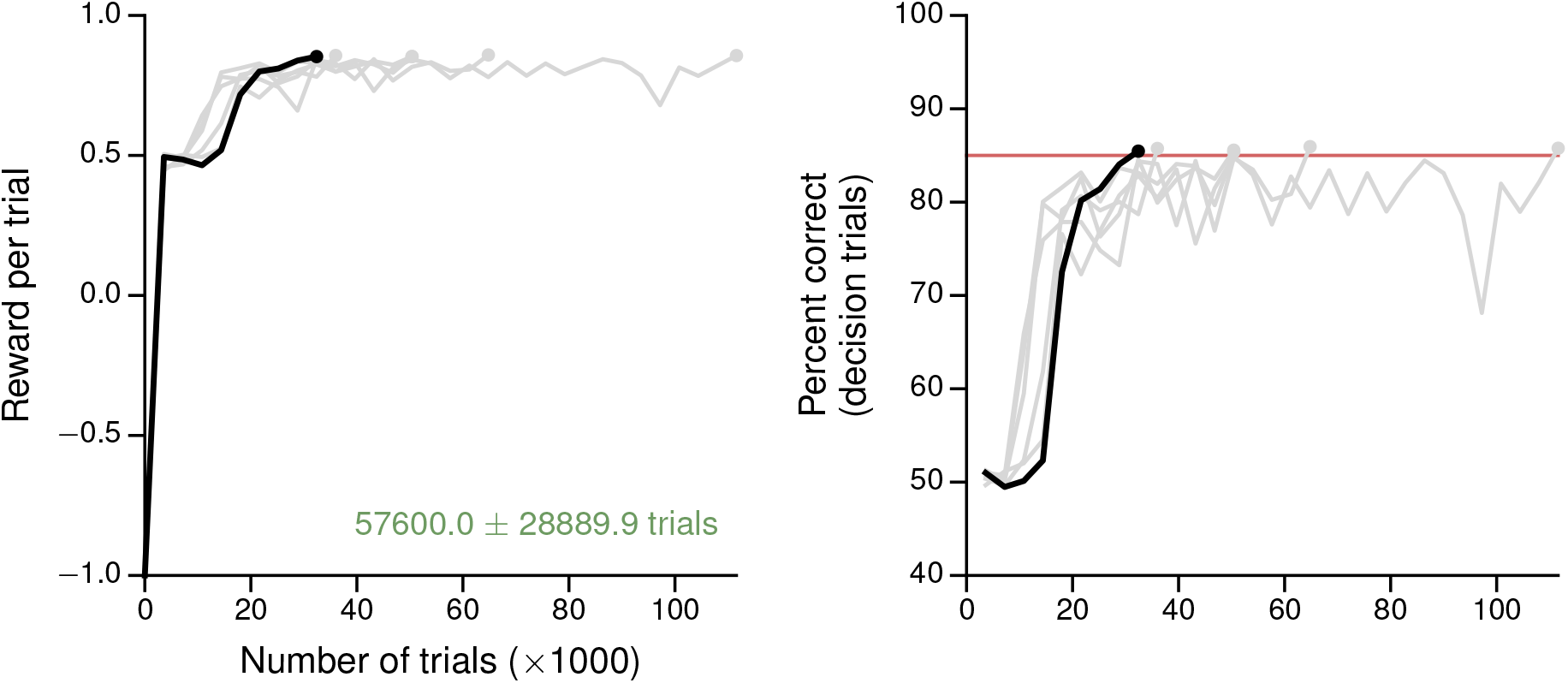
Figure 2—figure supplement 1. Learning curves for the context-dependent integration task. (**A**) Average reward per trial. Black is for the network realization in the main text, gray for additional realizations, i.e., trained with different random number generator seeds. (**B**) Percent correct, for trials on which the network made a decision (≥ 99% required for termination). Red: target performance (when training was terminated).

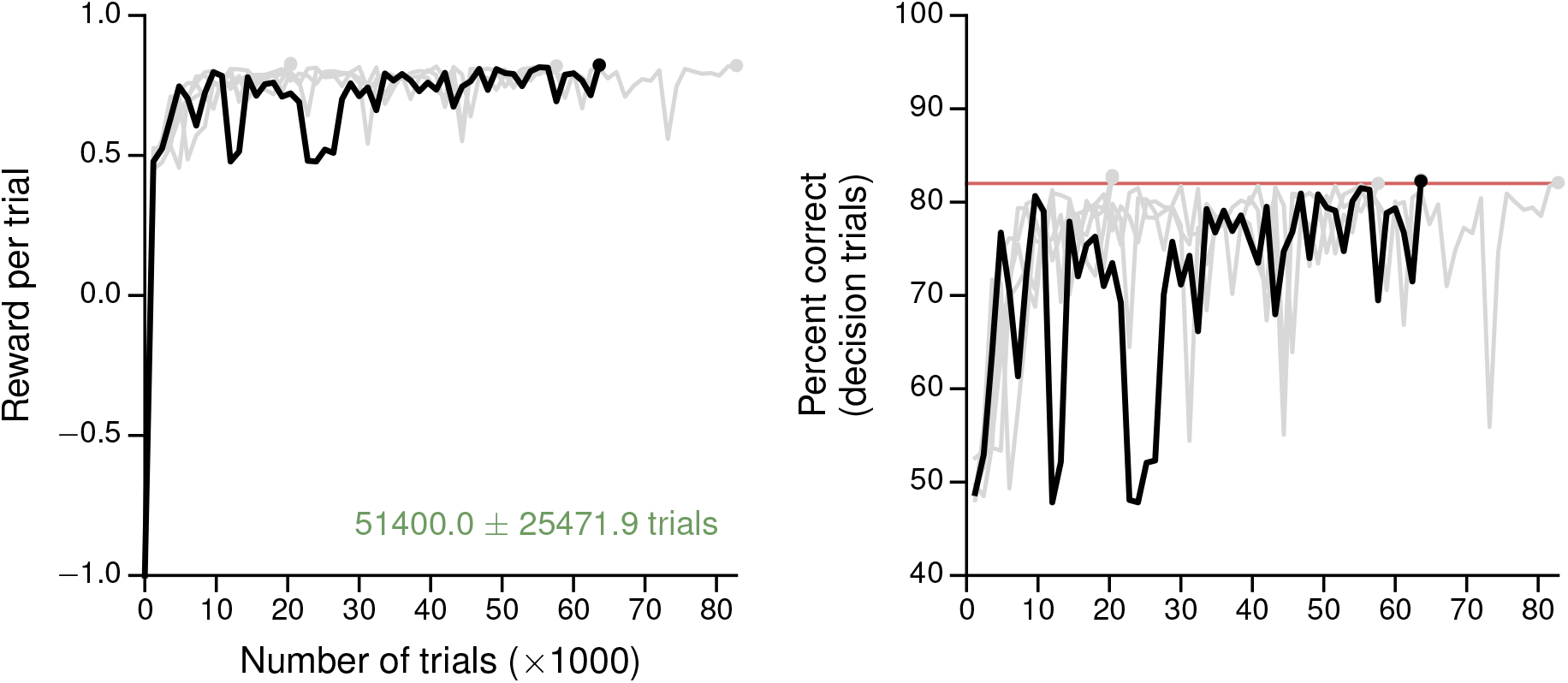
Figure 2—figure supplement 2. Learning curves for the multisensory integration task. (**A**) Average reward per trial. Black indicates the network realization shown in the main text, gray additional realizations, i.e., trained with different random number generator seeds. (**B**) Percent correct, for trials on which the network made a decision (≥ 99% required for termination). Red: target performance (when training was terminated).

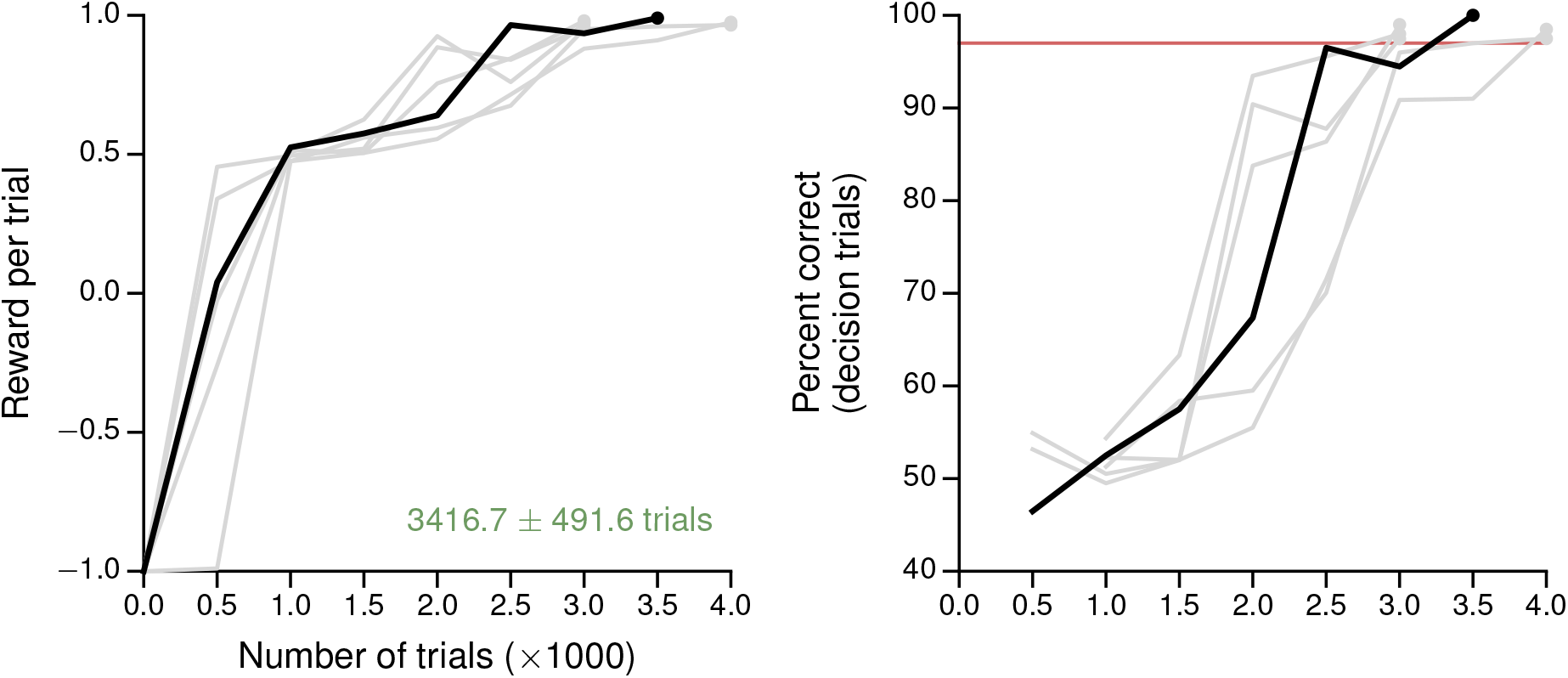
Figure 2—figure supplement 3. Learning curves for the parametric working memory task. (**A**) Average reward per trial. Black indicates the network realization shown in the main text, gray additional realizations, i.e., trained with different random number generator seeds. (**B**) Percent correct, for trials on which the network made a decision (≥ 99% required for termination). Red: target performance (when training was terminated).

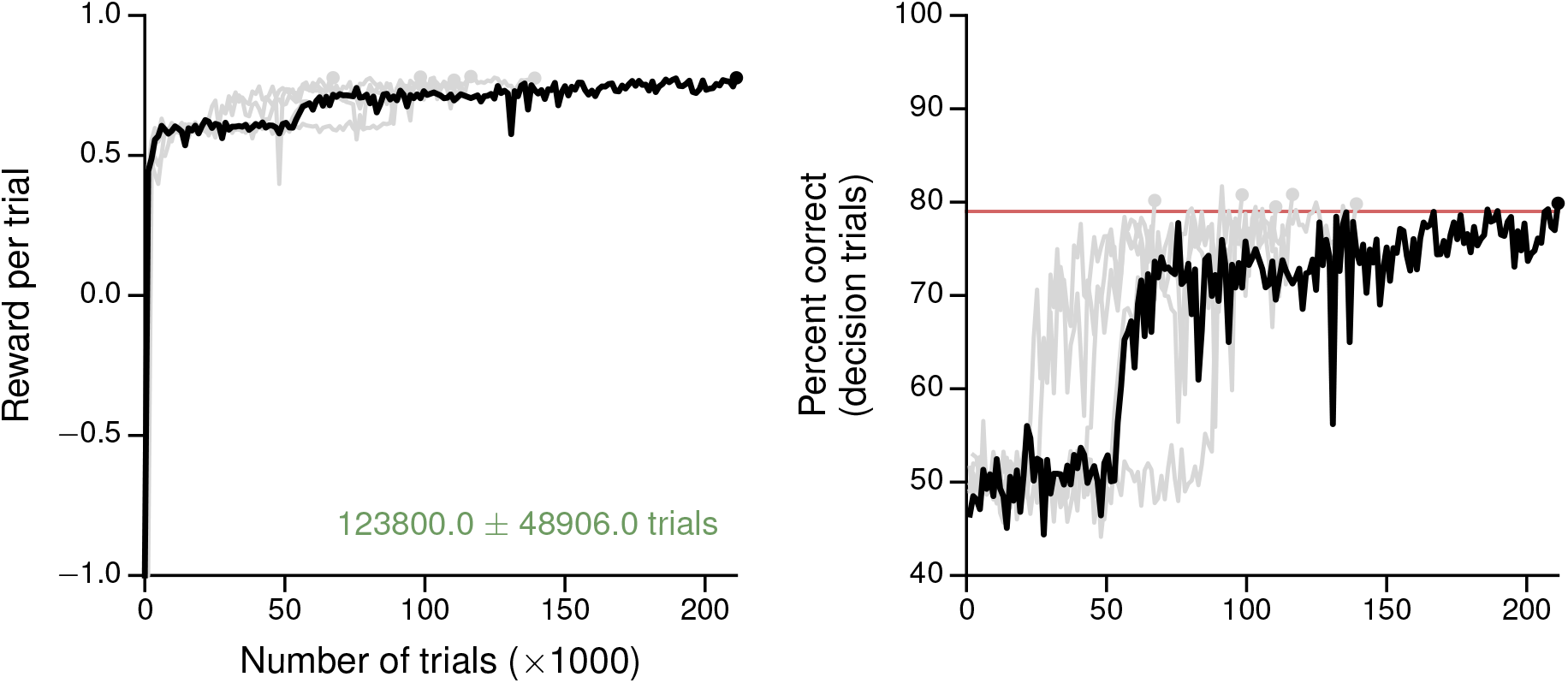
Figure 3—figure supplement 1. Learning curves for the postdecision wager task. (**A**) Average reward per trial. Black indicates the network realization shown in the main text, gray additional realizations, i.e., trained with different random number generator seeds. (B) Percent correct, for trials on which the network made a decision (≥ 99% required for termination). Red: target performance when the sure bet was accepted between 40-50% of the time.

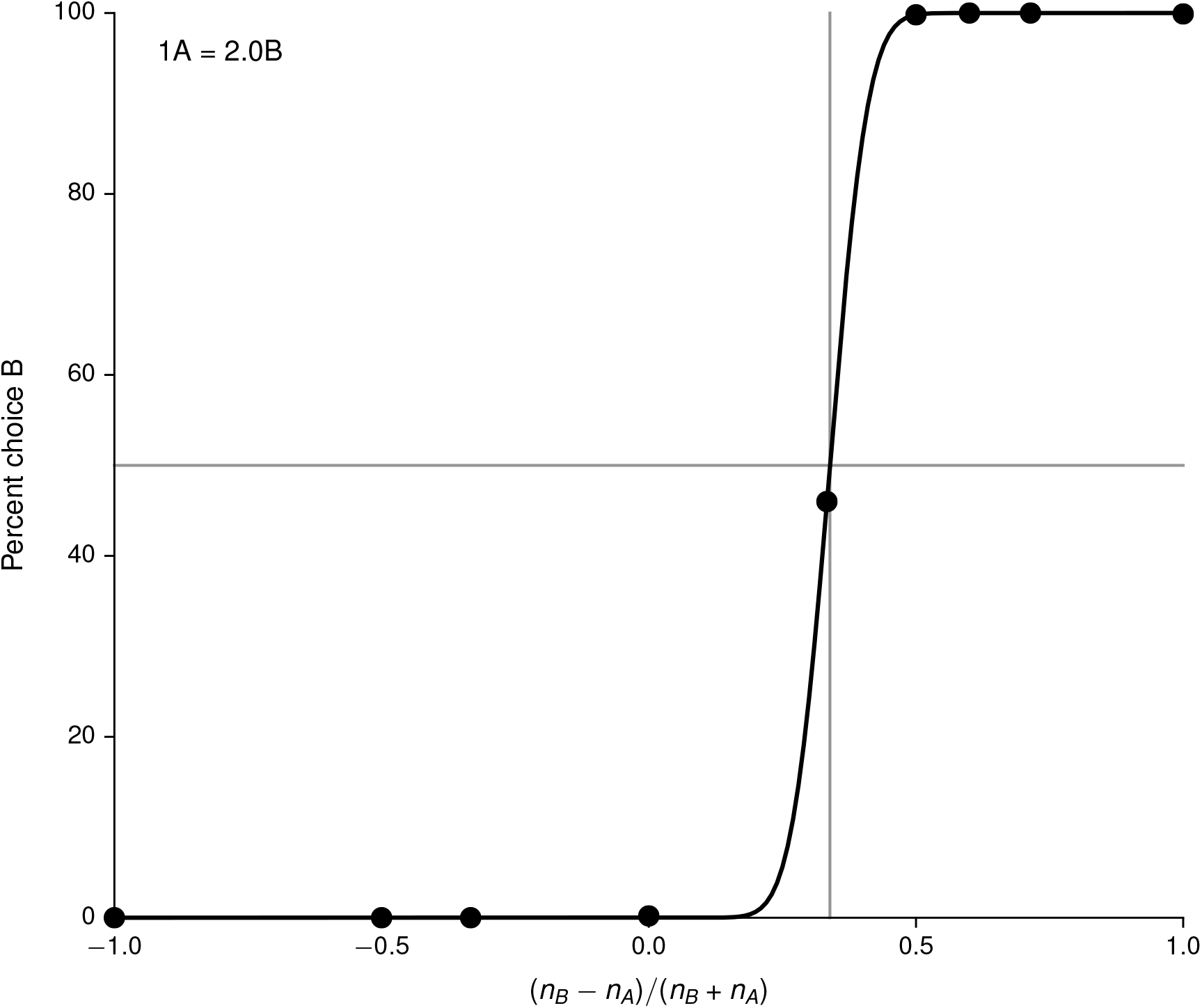
Figure 4—figure supplement 1. Fit of cumulative Gaussian with parameters *μ*, *σ* to the choice pattern in Figure 4 (upper), and the deduced indifference point 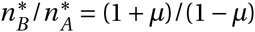

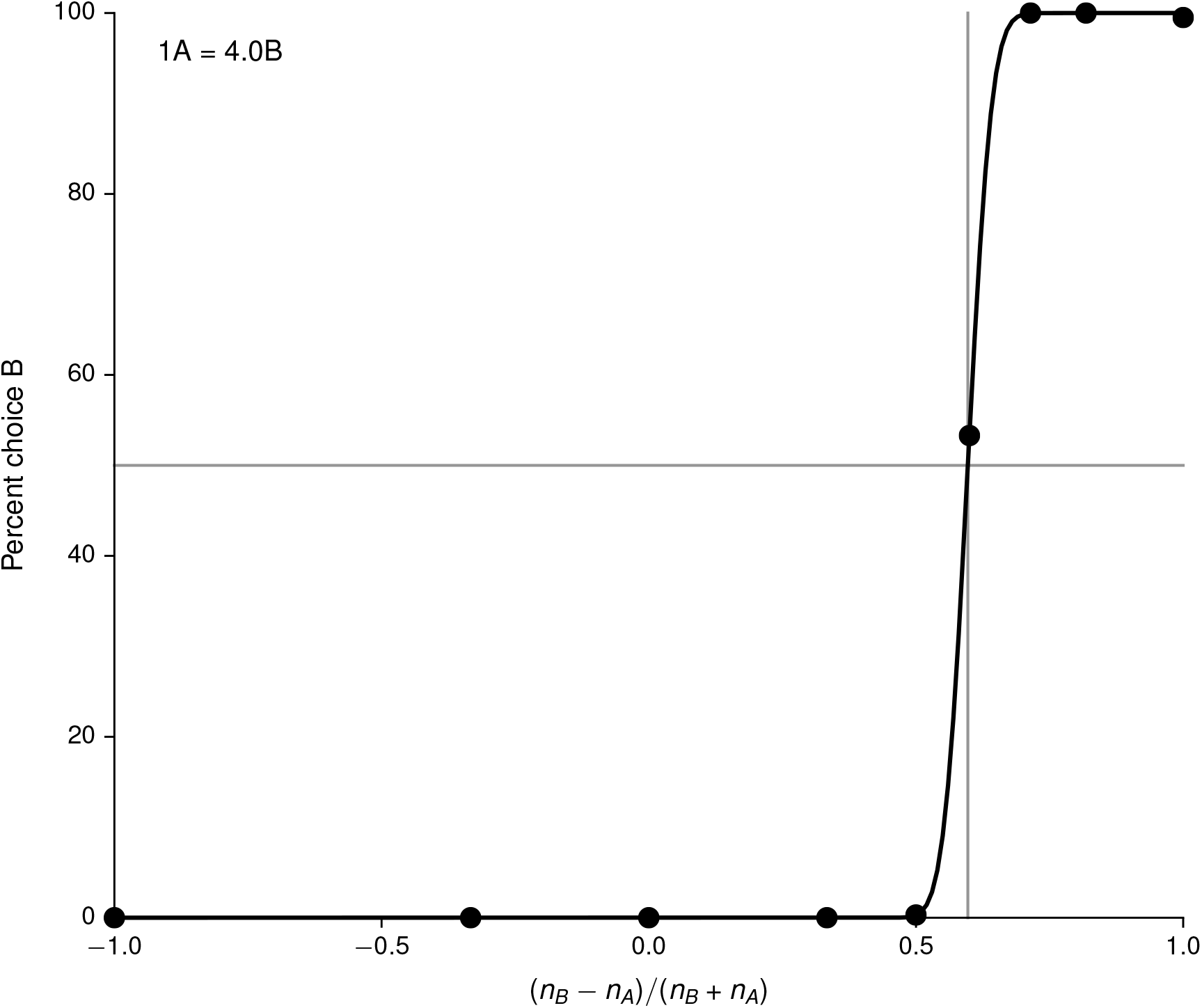
Figure 4—figure supplement 2. Fit of cumulative Gaussian with parameters *μ*, *σ* to the choice pattern in Figure 4A (lower), and the deduced indifference point 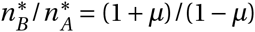.

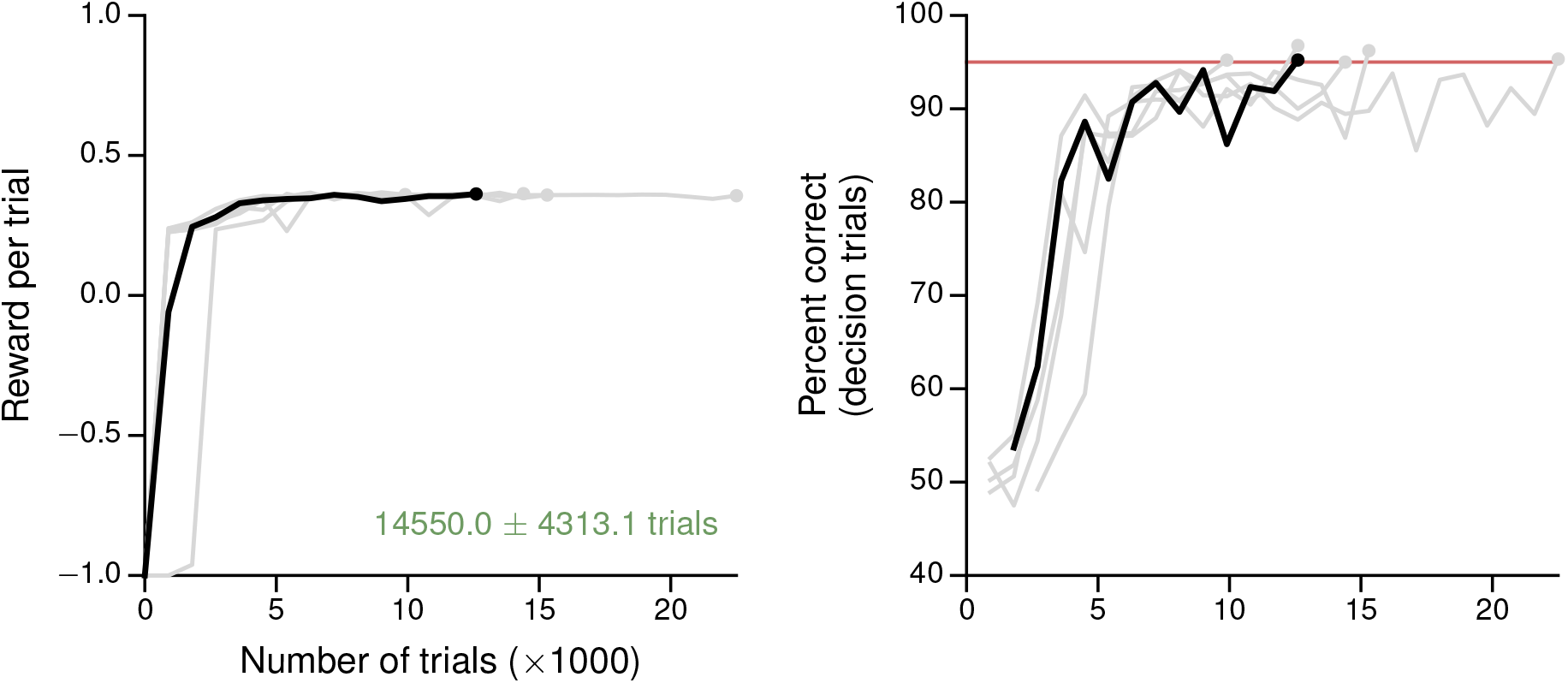
Figure 4—figure supplement 3. Learning curves for the value-based economic choice task. (**A**) Average reward per trial. Black indicates the network realization shown in the main text, gray additional realizations, i.e., trained with different random number generator seeds. (**B**) Percentage of trials on which the network chose the option that resulted in greater (or equal) reward, for trials where the network made a decision (≥ 99% required for termination). Note this is conceptually different from the previous tasks, where “correct” depends on the sensory inputs, not the rewards. Red: target performance (when training was terminated).

